# TLR priming licenses NAIP inflammasome activation by immunoevasive ligands

**DOI:** 10.1101/2023.05.04.539437

**Authors:** James P. Grayczyk, Marisa S. Egan, Luying Liu, Emily Aunins, Meghan A. Wynosky-Dolfi, Scott Canna, Andy J. Minn, Sunny Shin, Igor E. Brodsky

**Author notes:** Current address: Oncology Discovery, AbbVie Inc., 1 North Waukegan Rd., North Chicago, IL 60064. Current address: Immunology Research Unit, GlaxoSmithKline, Collegeville PA.

## Abstract

NLR family, apoptosis inhibitory proteins (NAIPs) detect bacterial flagellin and structurally related components of bacterial type III secretion systems (T3SS), and recruit NLR family, CARD domain containing protein 4 (NLRC4) and caspase-1 into an inflammasome complex that induces pyroptosis. NAIP/NLRC4 inflammasome assembly is initiated by the binding of a single NAIP to its cognate ligand, but a subset of bacterial flagellins or T3SS structural proteins are thought to evade NAIP/NLRC4 inflammasome sensing by not binding to their cognate NAIPs. Unlike other inflammasome components such as NLRP3, AIM2, or some NAIPs, NLRC4 is constitutively present in resting macrophages, and not thought to be regulated by inflammatory signals. Here, we demonstrate that Toll-like receptor (TLR) stimulation upregulates NLRC4 transcription and protein expression in murine macrophages, which licenses NAIP detection of evasive ligands. TLR-induced NLRC4 upregulation and NAIP detection of evasive ligands required p38 MAPK signaling. In contrast, TLR priming in human macrophages did not upregulate NLRC4 expression, and human macrophages remained unable to detect NAIP-evasive ligands even following priming. Critically, ectopic expression of either murine or human NLRC4 was sufficient to induce pyroptosis in response to immunoevasive NAIP ligands, indicating that increased levels of NLRC4 enable the NAIP/NLRC4 inflammasome to detect these normally evasive ligands. Altogether, our data reveal that TLR priming tunes the threshold for NAIP/NLRC4 inflammasome activation and enables inflammasome responses against immunoevasive or suboptimal NAIP ligands.

**Significance Statement:** Cytosolic receptors in the neuronal apoptosis inhibitor protein (NAIP) family detect bacterial flagellin and components of the type III secretion system (T3SS). NAIP binding to its cognate ligand engages the adaptor molecule NLRC4 to form NAIP/NLRC4 inflammasomes culminating in inflammatory cell death. However, some bacterial pathogens evade NAIP/NLRC4 inflammasome detection, thus bypassing a crucial barrier of the immune system. Here, we find that, in murine macrophages, TLR-dependent p38 MAPK signaling increases NLRC4 expression, thereby lowering the threshold for NAIP/NLRC4 inflammasome activation in response to immunoevasive NAIP ligands. Human macrophages were unable to undergo priming-induced upregulation of NLRC4 and could not detect immunoevasive NAIP ligands. These findings provide a new insight into species-specific regulation of the NAIP/NLRC4 inflammasome.

## Introduction

Pattern recognition receptors (PRRs) sense pathogen-associated molecular patterns (PAMPs) and signatures of pathogenesis during bacterial infection, and promote innate immune responses to clear infection (1-3). Many gram-negative bacterial pathogens encode type III secretion systems (T3SS) to deliver effector proteins into host cells and use the motility organelle flagella to promote infection. However, in addition to effector proteins, the T3SS of many pathogens translocate flagellin as well as structural components of the T3SS apparatus into the host cell cytosol (4). The NLR family apoptosis inhibitor proteins (NAIPs) bind to cytosolic flagellin and T3SS components, resulting in assembly of a multimeric protein complex known as the inflammasome that plays a crucial role in host defense against bacterial infections (5-7).

In mice, NAIP5 and NAIP6 detect cytosolic flagellin of bacterial pathogens such as *Salmonella*, *Legionella*, and *Pseudomonas*, while NAIP1 and NAIP2 recognize portions of the T3SS termed the needle and inner rod subunits, respectively (6-13). In contrast, humans encode a single NAIP that broadly recognizes all three classes of these ligands (14-17). Upon binding to their cognate ligands, NAIPs recruit the adaptor protein nucleotide-binding domain, leucine-rich repeat-containing protein caspase recruitment domain containing protein 4 (NLRC4), which in turn recruits and activates the zymogen protease caspase 1 (Casp1) within the inflammasome (14, 15, 18-23). Active Casp1 cleaves the pro-form of interleukin (IL)-1 family cytokines IL-1β and IL-18, as well as the pore-forming protein gasdermin D (GSDMD) to mediate pyroptosis (24-26).

The NAIP/NLRC4 inflammasome serves as an important barrier to bacterial colonization and infection. Notably, *Salmonella enterica* serovar Typhimurium (*S.* Tm) downregulates flagellin expression during its systemic phase, which is important for evasion of NLRC4-mediated host defense (27-31). Moreover, genetic resistance to *Legionella* infection in inbred mice is dependent on a functional *Naip5* gene, and *Legionella* expressing flagellin are rapidly cleared in a NAIP5/NLRC4-dependent manner (9, 12, 22). Additionally, wild-type mice are resistant to colonization by the primate-specific enteric pathogen, *Shigella flexneri*, whereas *Nlrc4^-/-^* mice are highly susceptible (32, 33).

NAIP5 binds to the flagellin D0 domain, which consists of N- and C-terminal alpha-helical regions (34, 35). Notably, the C-terminal 35 amino acids of the D0 domain are both necessary and sufficient for NAIP5/NLRC4 activation (36). Intriguingly, a small number of key residues at the C-terminus distinguish between closely related flagellins that activate or do not activate NAIP/NLRC4 inflammasomes. Specifically, activating flagellins from *S*.Tm, *Legionella*, or *Pseudomonas* all have a C-terminal arginine. In contrast, non-activating flagellins from *Helicobacter pylori*, Enteropathogenic *Escherichia coli* (EPEC), *Burkholderia thailandensis*, and several commensal bacteria lack the C-terminal arginine and instead contain C-terminal glutamine, glutamine-glycine, or threonine residues (7, 37, 38). These flagellins do not bind NAIP5 in yeast two-hybrid assays (7), implying that NAIP binding determines activation of the NAIP/NLRC4 inflammasome. Indeed, structural studies indicate that NAIP/NLRC4 inflammasome assembly is initiated by a single flagellin binding a single NAIP, leading to recruitment and polymerization of NLRC4 subunits and formation of the inflammasome (19, 21, 35, 37, 39).

In contrast to the sensor proteins NLRP3, AIM2, and NAIP1 (11, 40, 41), adaptor proteins such as NLRC4 are not widely considered to be transcriptionally induced in response to inflammatory signals, beyond a role for the transcription factors IRF8 and BRD4 in basal expression of *Nlrc4* and the *Naips* (42, 43). Whether inflammatory signals regulate NLRC4 expression and the potential consequences of such regulation on NAIP/NLRC4 inflammasome function is not known. Here, using a *Yersinia* T3SS-based system to deliver the D0 domain of flagellin or other NAIP ligands into macrophages, we find that TLR-induced priming enabled NAIP/NLRC4 detection of EPEC flagellin, which is normally not sensed in the absence of TLR priming. TLR stimulation upregulated NLRC4 expression, which correlated with enhanced NAIP/NLRC4 inflammasome-dependent responses. This enhanced NAIP/NLRC4 response required MyD88 and p38 MAPK signaling, as well as NAIP5. Intriguingly, TLR priming licensed NLRC4-dependent inflammasome responses to other evasive NAIP ligands, including the *S.* Tm SPI-2 inner rod and needle proteins, indicating that TLR priming lowers the threshold for NAIP/NLRC4 inflammasome activation in response to immunoevasive NAIP ligands. Surprisingly, we found that NLRC4 was not upregulated in TLR-primed human macrophages, correlating with lack of responsiveness to immunoevasive flagellins by human macrophages. Ectopic expression of murine or human NLRC4 in murine or human macrophages conferred enhanced inflammasome responses against immunoevasive flagellin, indicating that NLRC4 expression levels regulate responses to sub-optimal NAIP ligands. Our findings reveal that TLR priming licenses the NAIP-NLRC4 inflammasome detection of immunoevasive NAIP ligands by increasing NLRC4 expression. Furthermore, we uncover a key distinction between the ability of human and murine NAIP/NLRC4 inflammasomes to respond to immunoevasive NAIP ligands.

## Results

### C-terminal glutamine residues in flagellin disable recognition by the NAIP/NLRC4 inflammasome

Studies of the NAIP/NLRC4 inflammasome have used retroviral transduction, protein transfection, anthrax toxin-based delivery, and *L. monocytogenes* ActA-fusion proteins to deliver NAIP ligands into the host cell (10, 11, 13, 16, 17, 35, 36, 44). However, cytosolic delivery of flagellin, T3SS inner rod, or T3SS needle proteins under native conditions occurs via translocation through the T3SS or T4SS itself (4). We therefore sought to develop a T3SS-based method to deliver flagellin into the cell in order to study mechanisms of NAIP/NLRC4 inflammasome activation in response to T3SS-based delivery of NAIP ligands. Specifically, we developed a heterologous delivery system using the T3SS of the enteric gram-negative pathogen *Yersinia pseudotuberculosis* (*Yp*) by fusing the C-terminal FliCD0 region of flagellin to the N-terminal 100 amino acids of the T3SS-secreted *Yersinia* outer protein E (YopE^1-100^), which contains the YopE N-terminal secretion and translocation signal (**Fig. *S*1**). *Yp* represses expression of its own flagella during infection, allowing us to avoid any inflammatory responses by *Yp* flagella (45, 46). We also used *Yp* on a Δ*yopJ* background to prevent activation of YopJ-induced cell death (47-49). As expected, Δ*yopJ Yp* expressing *S.* Tm FliCD0 YopE fusion (FliCD0), but not EPEC FliCD0 YopE fusion (EPEC FliCD0) or the first 100 amino acids of YopE alone (YopE^1-100^), induced significant cell death of bone marrow-derived macrophages (BMDMs) in an NLRC4- and Casp1/11-dependent manner (**Fig. 1*A***). Consistently, delivery of *S.* Tm FliCD0 but not of EPEC FliCD0 or YopE^1-100^ induced secretion of IL-1β and cleavage of both Casp1 and GSDMD, which are markers of pyroptosis (**Fig. 1*B* and *C***).

**Figure 1.**
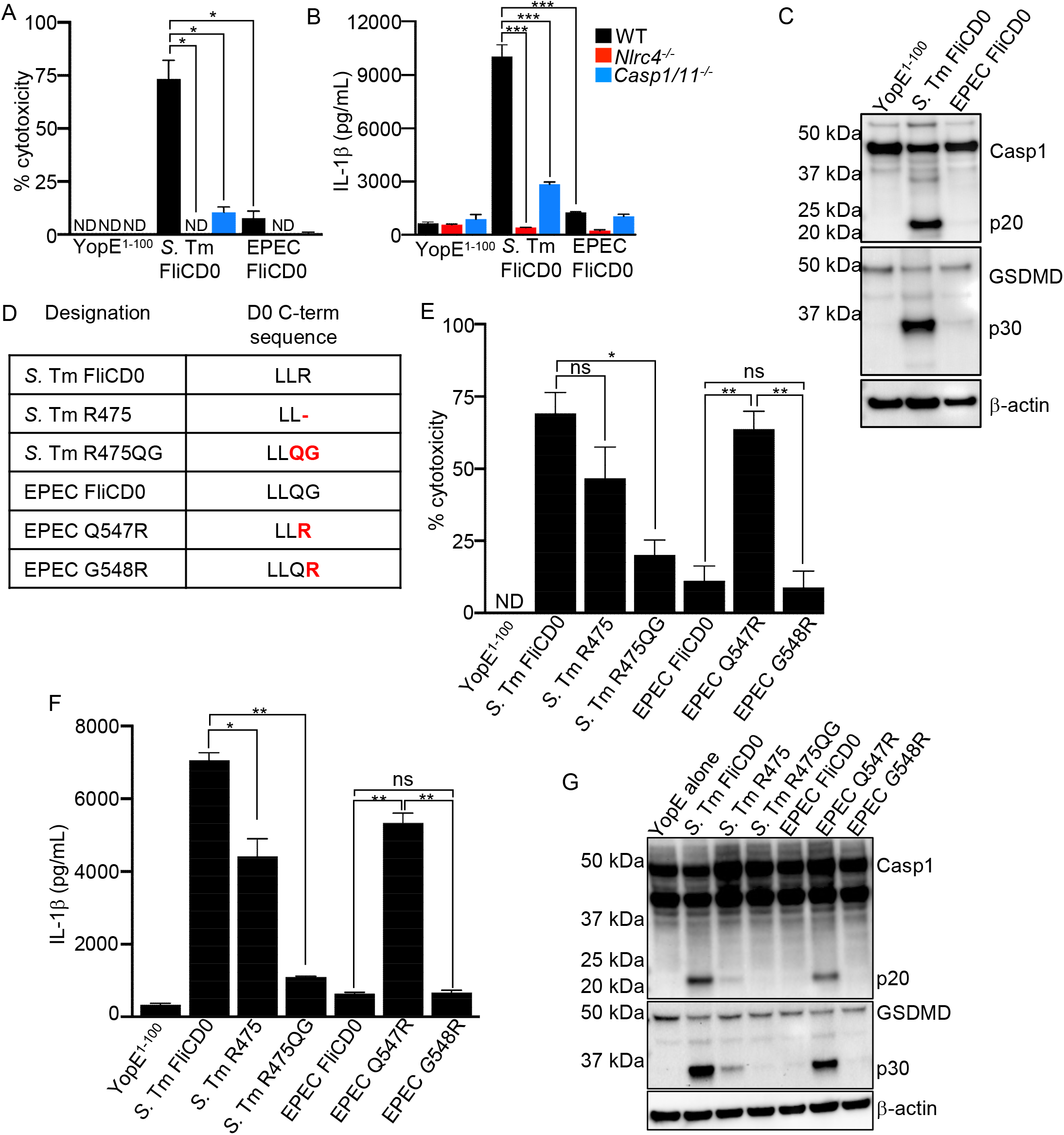
C-terminal glutamine residue abrogates NAIP inflammasome recognition of flagellin. (A-B) WT, *Nlrc4^-/-^* and *Casp1/11^-/-^* murine BMDMs were infected with Δ*yopJ Yp* expressing either YopE^1-100^ or YopE^1-100^ fused to the C-terminal D0 portion of FliC from *S.* Tm or EPEC for four hours at an MOI of 20. (A) % cytotoxicity was measured via lactate dehydrogenase (LDH) release. (B) IL-1β release (pg/mL) was measured by ELISA. (C) BMDMs were infected with the indicated strains for two hours at an MOI of 20. Combined supernatants and cellular lysates were analyzed by immunoblotting for Casp1, GSDMD, and β−actin (loading control). Representative of three independent experiments. (D) List of mutants generated to assess contribution of C-terminal residues in *S.* Tm or EPEC FliC D0. Red indicates the base pair mutations compared to the WT sequence. (E, F) WT murine BMDMs were infected with the indicated strains for four hours at an MOI of 20. (E) % cytotoxicity was measured via LDH release. (F) IL-1β release (pg/mL) was measured by ELISA. (G) BMDMs were infected with the indicated strains for one hour at an MOI of 20. Combined supernatants and whole cell lysates were analyzed by immunoblotting for Casp1, GSDMD, and β−actin (loading control). Representative of three independent experiments. (A, E) Data shown are the pooled means ± SEM from three independent experiments. Paired *t* test was performed to assess statistical significance. (B, F) Data shown are the pooled means ± SEM from three independent experiments. Statistical significance was measured by performing an unpaired *t* test. ND=not detected; ns=not significant; *,*P*<0.05, **,*P*<0.01,***,*P*<0.001.

Three conserved leucine residues at the C-terminus of flagellin confer binding by NAIP5 and subsequent activation the NAIP5/NLRC4 inflammasome (36). However, these leucines are retained in flagellins from bacteria such as *E. coli* that do not activate the inflammasome, indicating that other residues are also important for NAIP5 binding. Recent studies employing ectopic expression or yeast two-hybrid studies indicate bacterial flagellins that contain a C-terminal arginine, including *Salmonella*, *Legionella*, and *Pseudomonas,* activate NAIP5 (7, 37, 38). In contrast, flagellins that contain a C-terminal glutamine, glutamine-glycine, or residue other than arginine, including *E. coli*, *Burkholderia*, or *Helicobacter* do not bind NAIP5 or activate the NAIP5/NLRC4 inflammasome (7, 37, 38). Multiple mutations in flagellin that also impact motility are proposed to be necessary to evade NAIP5 recognition (35), making it surprising that a single terminal amino acid would have such an impact on activation of NAIP5/NLRC4. We therefore sought to test the requirement for the C-terminal arginine in NAIP5/NLRC4 inflammasome activation under conditions where flagellin was delivered via a T3SS. We generated an arrayed series of isogenic constructs in which the terminal residue(s) of *S*.Tm or EPEC FliCD0 were mutated to swap glutamine-arginine residues (**Fig. 1*D***). *S.* Tm FliCD0 with glutamine-glycine in place of arginine to mimic the terminal EPEC FliCD0 sequence led to significantly decreased inflammasome activation relative to WT *S.* Tm FliCD0 as assessed by cell death, IL-1β release, Casp1, and GSDMD cleavage (**Fig. 1*E*-*G***). Removing the terminal arginine of *S.* Tm FliCD0 also significantly reduced cytotoxicity and IL-1β release, and abrogated Casp1 and GSDMD cleavage, indicating that the C-terminal arginine is required for optimal NAIP5/NLRC4 inflammasome activation in response to flagellin (**Fig. 1*E*-*G***). Conversely, replacement of the EPEC FliCD0 C-terminal glutamine-glycine with arginine resulted in robust cell death, IL-1β release, as well as GSDMD and Casp1 cleavage at similar levels to S. Tm FliCD0 (**Fig. 1*E*-*G***). Intriguingly, a glutamine at the C-terminus abrogated inflammasome activation even if the terminal amino acid itself was arginine, indicating that glutamine at the FliC D0 C-terminus acts as a dominant negative to suppress NAIP/NLRC4 inflammasome activation, whereas in the absence of glutamine, arginine is necessary and sufficient for NAIP/NLRC4 inflammasome activation. Together, these data demonstrate that the *Yp* T3SS delivery system recapitulates the currently known features of NAIP/NLRC4 inflammasome activation by flagellin, and demonstrates a key role for the flagellin C-terminal amino acid in NAIP5 sensing.

### TLR priming licenses NAIP/NLRC4 inflammasome activation by immunoevasive NAIP ligands

PRR signaling during bacterial infection upregulates absent in melanoma (AIM2)-like receptors and NLRP3 to facilitate detection of intracellular PAMPs or DAMPs and is necessary for activation of these inflammasomes (40, 41). While *Naip1* in murine BMDMs is transcriptionally induced by poly-I:C and LPS stimulation to enable inflammasome activation by the bacterial T3SS needle (11), *Nlrc4* is not currently known to be transcriptionally upregulated in response to PAMP stimulation. However, immune cells at mucosal surfaces are exposed to numerous PAMP-PRR signals from commensal microbes, and infection provides additional exogenous and endogenous inflammatory mediators. We therefore considered the possibility that priming of the NAIP/NLRC4 inflammasome enhances NAIP sensitivity so that evasive ligands like EPEC FliCD0 can be sensed. To test this, we primed murine BMDMs with synthetic triacylated lipopeptide (Pam3CSK4 or Pam3), diacylated lipopeptide (Pam2CSK4 or Pam2), LPS, or the inflammatory cytokine IFN-γ. Intriguingly, BMDMs primed with the TLR ligands Pam3, Pam2, or LPS, but not IFN-γ, exhibited increased cell death and IL-1β release in response to EPEC FliCD0 (**Fig. *S*2*A,B***). Cell death and IL-1β release in Pam3-primed cells were dependent on NLRC4, Casp1, and NAIP5, indicating that priming enhances the ability of the NAIP5/NLRC4 inflammasome to detect immunoevasive flagellins, rather than enabling NAIP6 to compensate for NAIP5 (**Fig. 2*A,B***). As expected, this increased cytotoxicity and IL-1β release in Pam3-primed cells in response to EPEC FliCD0 correlated with elevated GSDMD cleavage and Casp1 cleavage (**Fig. 2*C***). Notably, Pam3 priming also induced elevated cytotoxicity and IL-1β release in response to mutant versions of *S*. Tm FliCD0 that contain C-terminal glutamine, indicating that TLR priming overcomes the inhibitory effect of glutamine on NAIP5/NLRC4 activation (**Fig. S3*A,B***). In Pam3-primed cells, control infection with *Yp* expressing YopE^1-100^ alone induced significant levels of IL-1β release with minimal levels of cytotoxicity or processing of Casp1 or GSDMD (**Fig. 2*A-C***). However, this release of IL-1β in response to YopE alone was independent of NAIP5, indicating that this response is distinct from the NAIP/NLRC4-depenent pathway induced by flagellin, but was still NLRC4 and Casp1/11-dependent, suggesting that it was likely due to priming-induced responses to *Yersinia* T3SS needle or rod proteins (50) (**Fig. 2*B***).

**Figure 2.**
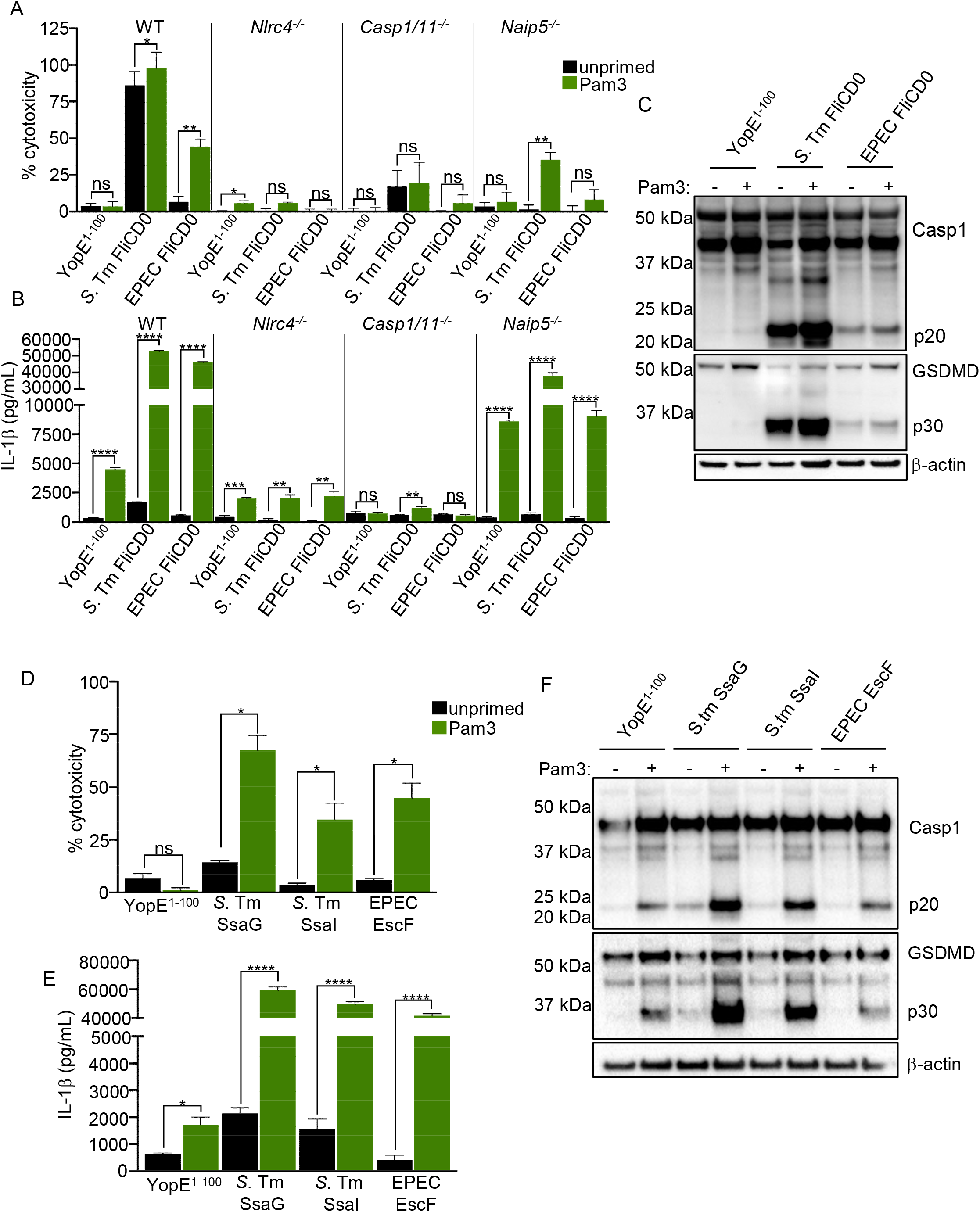
TLR priming enables murine macrophages to recognize immunoevasive NAIP ligands. (A and B) WT, *Nlrc4^-/-^*, *Casp1/11^-/-^*, and *Naip5^-/-^* murine unprimed BMDMs or primed with 0.5 μg/mL Pam3CSK4 (Pam3) for 16 hours were infected with the indicated strains for four hours at an MOI of 20. (A) Cell death (% cytotoxicity) was measured via lactase dehydrogenase (LDH) release. (B) IL-1β release (pg/mL) was measured by ELISA. (C) BMDMs were infected with the indicated strains for two hours at an MOI of 20, TCA-precipitated supernatants and whole cell lysates were collected and combined. Samples were analyzed by immunoblot for Casp1, GSDMD, and β−actin (loading control). Image representative of three independent experiments. (D andE) WT murine unprimed BMDMs or primed with 0.5 μg/mL Pam3CSK4 for 16 hours were infected with the indicated strains for two hours at an MOI of 20. (D) % cytotoxicity was measured via LDH release. (E) IL-1β release (pg/mL) was measured by ELISA. (F) BMDMS were infected with the indicated strains for four hours at an MOI of 20. Combined supernatants and whole cell lysates were analyzed by immunoblotting for Casp1, GSDMD, and β−actin (loading control). Image representative of three independent experiments. (A and D) Data shown are pooled means ± SEM from three independent experiments. A paired *t* test was performed to assess statistical significance. (B, G) Data shown are representative of three independent experiments and are the combined means ± SEM. Statistical significance was measured by performing an unpaired *t* test. ns=not significant; *, *P*<0.05, **, *P*<0.01,***, *P*<0.001,****, *P*<0.001.

The NLRP3 inflammasome requires priming (termed ‘Signal 1’) to undergo inflammasome activation (40, 51), and can synergize with NLRC4 for maximal responses to *S*. Tm and other bacterial pathogens (16, 52, 53). Notably, *Nlrp3^-/-^* BMDMs were unaffected in their ability to undergo priming-induced cytotoxicity or IL-1β release in response to cytosolic delivery of EPEC FliCD0 (**Fig. S3*C,D***), indicating that NLRP3 does not contribute to TLR-induced priming of NAIP/NLRC4 responses to immunoevasive flagellin. This was also the case for BMDMs lacking the inflammasome adaptor protein ASC, indicating that TLR-induced activation of the NAIP/NLRC4 inflammasome by evasive EPEC FliCD0 requires an intact NAIP-NLRC4-Casp1 signaling axis but not NLRP3 or ASC to drive pyroptosis (**Fig. S3*E,F***).

In addition to NAIP5/NLRC4 inflammasome evasion by changes in the C-terminal FliCD0 domain, a number of pathogens produce T3SS needle and inner rod proteins that also evade NAIP/NLRC4 detection in murine macrophages to enable intracellular replication (10). This includes the SsaI inner rod subunit of the SPI-2 T3SS in *S*. Tm and the SPI-2 T3SS needle protein SsaG, though human NAIP is capable of detecting (16). Furthermore, the EPEC needle, EscF (34, 54), is also not sensed by the NAIP/NLRC4 inflammasome (8), raising the question of whether TLR priming of the NAIP/NLRC4 inflammasome extends to recognition of other classes of immunoevasive NAIP ligands in addition to flagellin. We therefore evaluated cytotoxicity of Pam3-primed and unprimed BMDMs infected with *Yp* expressing YopE fused to *S*. Tm SsaG or SsaI, as well as EPEC EscF. As expected, unprimed BMDMs did not respond to these ligands (**Fig. 2*D,E***). However, Pam3-primed BMDMs exhibited significantly increased cell death and IL-1β release in response to SsaG, SsaI, and EscF, suggesting that murine macrophages are able to detect immunoevasive NAIP ligands when stimulated through TLR2 (**Fig. 2*D,E***). Similar to SPI-2 ligands, priming of BMDMs lead to increased cell death and IL-1β release in response to cytosolic delivery of EscF (**Fig. 2*D,E***). The increase in cell death and IL-1β release in primed cells in response to T3SS components correlated with an increase in Casp1 and GSDMD cleavage (**Fig. 2F**). Altogether, our data reveal that TLR priming licenses the NAIP/NLRC4 inflammasome to detect normally evasive ligands.

### TLR priming upregulates NLRC4 expression to enable NAIP/NLRC4 inflammasome activation by evasive ligands

Pam3 and Pam2 are ligands for TLR2, which requires the adaptor MyD88 to transduce responses (55). Consistently, increased cell death in response to EPEC FliCD0 required MyD88 in BMDMs primed with Pam3 or Pam2, but not with LPS, suggesting that TRIF signaling can also induce this priming (**Fig. *S2A,B***). We next considered the possibility that TLR priming might impact the expression of *Naip* or *Nlrc4* genes. While *Naip1* is induced by poly(I:C), and steady-state levels of *Nlrc4* and *Naips* require the transcription factor IRF8 and epigenetic reader BRD4 (11, 42, 43), it is not thought the components of the NAIP/NLRC4 inflammasome are under inducible transcriptional control. Intriguingly, while we observed no increase in mRNA expression of *Naip1* or *2*, and a small decrease in expression of *Naip5* and *6*, *Nlrc4* levels exhibited a four-fold increase (**Fig. 3*A***). Furthermore, Pam3 signaling induced NLRC4 protein expression in a dose dependent manner (**Fig. 3*B***). Interestingly, we observed a trend toward increased transcript and protein levels at 8 hours post-priming, while the highest levels of *Nlrc4* transcript and NLRC4 protein occurred 16 hours post priming (**Fig. *S*4*A,B***). Notably, stimulation with either Pam2 or LPS, but not IFN-γ, also increased *Nlrc4* mRNA abundance and protein levels after 16 hours post priming (**Fig. *S*4*A,B***). These data indicate that TLR2/4 signaling, but not IFN-γ signaling, regulate NLRC4 expression in BMDMs.

**Figure 3.**
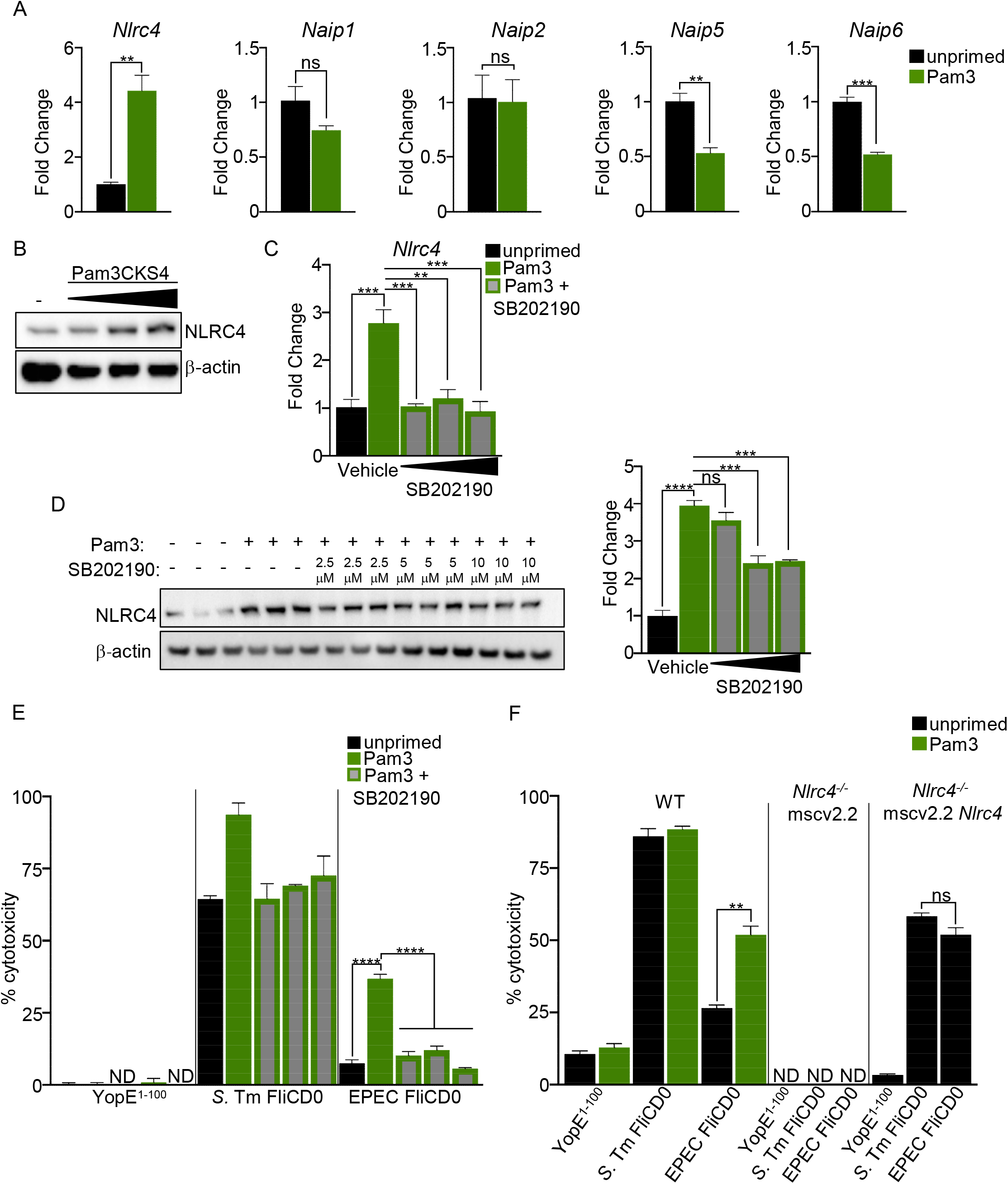
TLR priming through p38 MAPK signaling increases NLRC4 expression to promote NAIP/NLRC4 inflammasome responses to immunoevasive ligands. (A, B) WT murine BMDMs were unprimed or primed for 16hrs with (A) 0.5 μg/mL Pam3CSK4 or (B) 0.25 μg/mL, 0.5 μg/mL, or 1 μg/mL Pam3CSK4. *Nlrc4, Naip1, Naip2, Naip5,* and *Naip6* transcripts were assessed via qRT-PCR (A) or NLRC4 protein levels via immunoblot (B). Fold change was assessed relative to unstimulated BMDMs. Data representative of three independent experiments. (C-E) WT murine BMDMs were unprimed or primed with 0.5 μg/mL Pam3CSK4. One hour prior to priming, the BMDMs were treated with 2.5 μM, 5 μM, or 10 μM p38 MAPK inhibitor SB202190 or DMSO vehicle control. (C) *Nlrc4* transcript levels relative to *Gapdh* were assessed via qRT-PCR. Fold change was determined relative to unstimulated control. (D) NLRC4 protein levels were analyzed via immunoblot. NLRC4 protein intensity was quantified relative to loading controls, and average fold change ± SEM in NLRC4 protein levels was assessed relative to unstimulated DMSO-treated cells. Data representative of three independent experiments. (E) % cytotoxicity was measured via LDH release two hours post-infection with the indicated strains at an MOI of 20. (F) Immortalized ER-Hoxb8 *Nlrc4^-/-^* murine macrophages were transduced with the retroviral mscv2.2 vector alone (mscv2.2) or mscv2.2 overexpressing *Nlrc4* (mscv2.2-*Nlrc4*). WT macrophages, *Nlrc4^-/-^* macrophages transduced with mscv2.2, or *Nlrc4^-/-^* macrophages transduced with mscv2.2 *Nlrc4* were infected with indicated *Yp* strains expressing YopE^1-100^, *S.* Tm FliCD0, and EPEC FliCD0 for two hours at an MOI of 20. % cytotoxicity was measured via LDH release. Only WT ER-Hoxb8 cells were primed for 16 hours with Pam3CSK4. (A, C, E, F) Data shown are representative of three experiments. Means ± SEM are shown (n=3) and statistical significance was measured by performing an unpaired *t* test (A, F) or 1-way ANOVA (C-E). ND=not detected; ns=not significant; **, *P*<0.01,***, *P*<0.001,****, *P*<0.0001

We next sought to determine the basis for transcriptional induction of *Nlrc4* in response to TLR priming. TLR signaling leads to activation of both nuclear factor kappa B (NF-κB) and the mitogen activated protein kinase (MAPK) signaling cascades to promote inflammatory gene transcription. Recently, the MAPK p38 has been implicated in controlling NLRP1 inflammasome activation in response to cellular stress (56, 57). Furthermore, as BRD4 has been implicated in IRF8-mediated basal transcription of *Nlrc4* (42) and p38 can activate BRD4 activity in other cell types (58), we considered that p38 MAPK may contribute to TLR-based priming of the NAIP/NLRC4 inflammasome to immunoevasive NAIP ligands. Indeed, inhibition of p38 MAPK activity prevented Pam3-induced upregulation of *Nlrc4* transcript and protein (**Fig. 3*C,D***) as well as a known TLR-induced transcript, *Il6* (**Fig. S4*C***), but did not impact basal NLRC4 expression in the absence of priming (**Fig. S4*D,E***). Critically, p38 MAPK inhibition also abrogated Pam3-induced cytotoxicity in response to EPEC FliCD0, but not in response to *S*. Tm FliC D0 (**Fig. 3*E***). These data indicate that TLR stimulation upregulates NLRC4 via a p38 MAPK-dependent pathway and suggest that increased NLRC4 levels enable activation of the murine NAIP/NLRC4 inflammasome in response to suboptimal or immunoevasive ligands. To test whether increasing the expression of NLRC4 is sufficient to enable responses to immunoevasive flagellins, we ectopically expressed murine NLRC4 in immortalized *Nlrc4^-/-^* BMDMs (**Fig. S4*F***). Intriguingly, whereas WT iBMDMs exhibited Pam3 priming-dependent cytotoxicity in response to cytosolic delivery of EPEC FliCD0, iBMDMs overexpressing NLRC4 did not require priming in order to respond to EPEC FliCD0 (**Fig. 3*F***), demonstrating that increased NLRC4 expression is sufficient to enable NAIP/NLRC4 inflammasome detection of immunoevasive flagellins.

### Human NAIP/NLRC4 inflammasome cannot be primed to detect evasive flagellin ligands

In contrast to mice, humans express a single NAIP that broadly detects bacterial flagellin and T3SS structural components (14, 15, 17). Human NAIP is capable of detecting the SPI-2 needle protein SsaG from *Salmonella* to restrict intracellular infection, suggesting that the broader specificity of the single human NAIP could potentially enable detection of a wider range of ligands (16). As expected, human THP-1 monocyte-derived macrophages exhibited robust responses to *S*. Tm FliCD0 delivered by the *Yp* T3SS that was dependent on NLRC4 and NAIP (**Fig. 4*A***). Surprisingly however, THP-1 monocyte-derived macrophages did not respond to the EPEC FliCD0 domain, despite being basally stimulated with Pam3 (**Fig. 4*A***). Notably, and in marked contrast to their murine counterparts, THP-1 monocyte-derived macrophages exhibited reduced levels of *NLRC4* transcript when treated with Pam3, Pam2, or LPS, indicating that human and murine *NLRC4* genes exhibit distinct transcriptional responses to TLR stimulation (**Fig. 4*B***). Human *NAIP* transcript levels were similarly reduced in response to TLR stimuli, and contrasted with the control gene *IL6*, which was dramatically upregulated by all stimuli (**Fig. 4*B***). Moreover, in contrast to murine NLRC4, human NLRC4 protein levels were unaltered in response to Pam3 stimulation, suggesting that the human NAIP/NLRC4 inflammasome does not undergo priming in response to TLR ligands (**Fig. 4*C***). Importantly, primary human monocyte-derived macrophages also failed to detect the EPEC FliCD0 domain even after Pam3 priming, as levels of IL-1β secreted in response to EPEC FliCD0 were identical to the YopE1-100 alone, in contrast to *S*. Tm FliCD0 (**Fig. 4*D***). Similarly to murine NLRC4, expression of human NLRC4 in immortalized murine BMDMs resulted in priming-independent responses to cytosolic delivery of EPEC FliCD0 (**Fig. 4*E***). Critically, overexpression of human NLRC4 in human THP-1 monocyte-derived macrophages was also sufficient to enable human cells to detect the EPEC FliCD0 domain (**Fig. 4*F***). Together these data indicate that in contrast to murine macrophages, human NLRC4 is deficient in its ability to undergo TLR-induced priming, and that expression levels of NLRC4 also modulate the ability of the human NAIP/NLRC4 inflammasome to undergo activation in response to evasive ligands. Overall, our findings reveal an unexpected role for NLRC4 expression in tuning the responsiveness of the NAIP/NLRC4 inflammasome to immunoevasive ligands.

**Figure 4.**
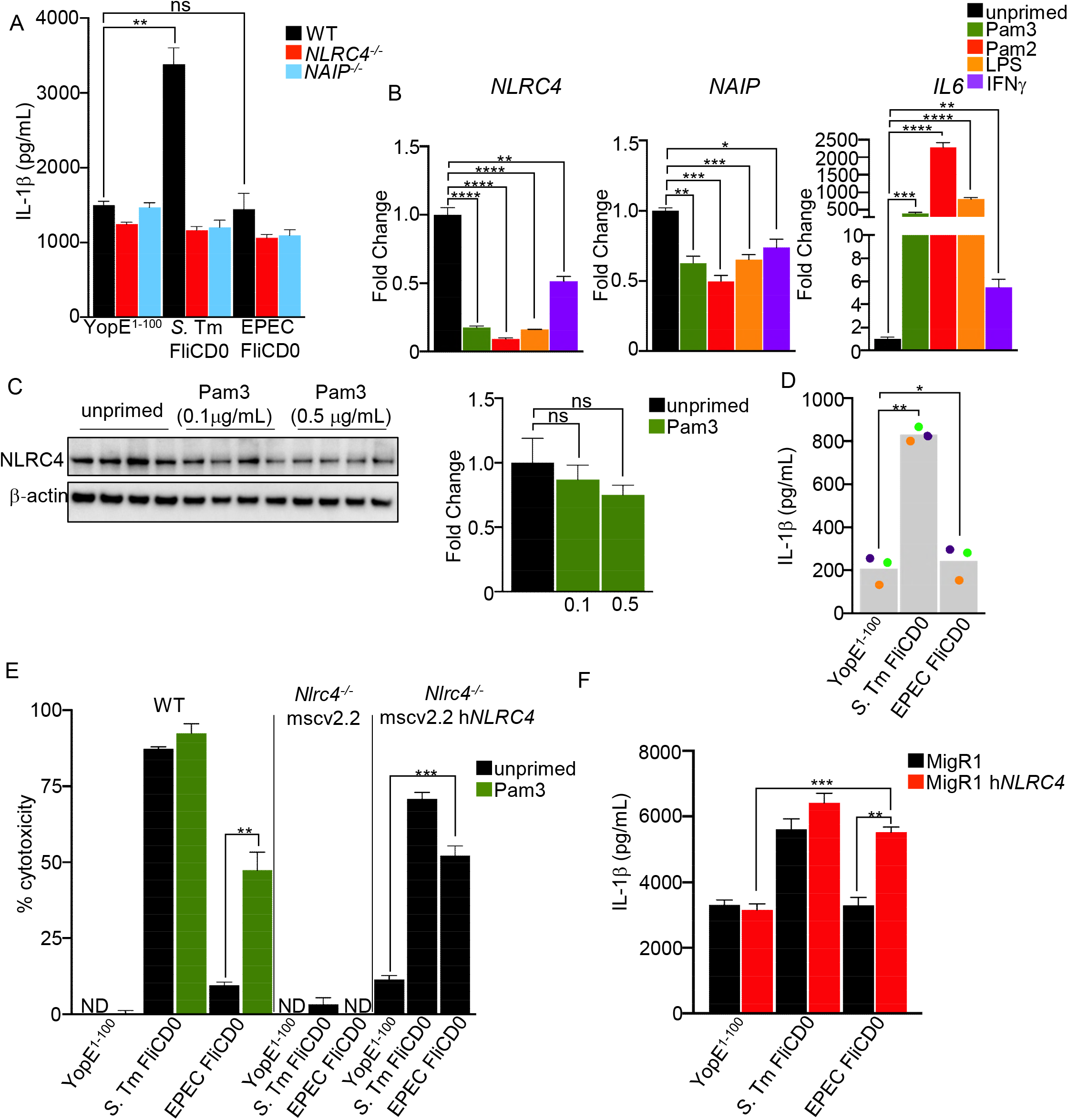
The human NAIP/NLRC4 inflammasome does not respond to TLR priming. (A) WT, *Nlrc4^-/-^*, or *Naip^-/-^* THP-1 macrophages were primed with 0.1 μg/mL Pam3CSK4 for 16 hours and infected with WT *Yp* expressing either YopE^1-100^, *S.* Tm FliCD0, or EPEC FliCD0 at an MOI of 60 for six hours. IL-1β release (pg/mL) was measured by ELISA. (B, C) WT THP-1 macrophages were primed with 0.1 μg/mL (B, C) or 0.5 μg/mL (C) Pam3CSK4, 0.05 μg/mL Pam2CSK4, 100 ng/mL LPS, or 100 ng recombinant human IFN-γ for 16 hours. (B) *Nlrc4, Naip,* and *Il6* transcript levels relative to *Hprt* were assessed via qRT-PCR. Fold change in transcript levels was analyzed relative to unstimulated control. (C) NLRC4 protein levels were analyzed via immunoblot. NLRC4 protein intensity was quantified relative to loading controls, and fold change in NLRC4 protein levels was assessed relative to unstimulated cells. Data representative of three independent experiments. (D) Primary hMDMs from three healthy human donors were primed with 0.1 μg/mL Pam3CSK4 for 16 hours and infected with indicated strains at an MOI of 60 for 6 hours. IL-1β release (pg/mL) was measured by ELISA. Each dot represents the mean of each donor derived from triplicate wells. Bars depict the mean of three donors. (E) Immortalized ER-Hoxb8 *Nlrc4^-/-^* murine macrophages were transduced with retroviral mscv2.2 overexpressing human *NLRC4* (mscv2.2-h*NLRC4*). WT, *Nlrc4^-/-^* mscv2.2, or *Nlrc4^-/-^* mscv2.2 h*NLRC4* macrophages were infected with indicated strains for two hours at an MOI of 20. % cytotoxicitywas measured via LDH release. Only WT Hoxb8 cells were primed for 16 hours with Pam3CSK4. (F) THP-1 human monocyte-derived macrophages transduced with the MigR1 retroviral vector alone or overexpressing h*NLRC4* were primed with 0.1 μg/mL Pam3CSK4 for 16 hours and infected with WT *Yp* expressing either YopE^1-100^, *S.* Tm FliCD0, or EPEC FliCD0 at an MOI of 60 for six hours. IL-1β release (pg/mL) was measured by ELISA. (A-F) Data shown are representative of three independent experiments. Means ± SEM are shown (n=3) and statistical significance was measured by performing an unpaired *t* test (A, C, E, F) or paired *t* test (D) or 1-way ANOVA (B). ND=not detected; ns=not significant; *, *P*<0.05, **, *P*<0.01,***, *P*<0.001,****, *P*<0.0001.

### TLR priming overcomes evasion of the NAIP/NLRC4 inflammasome during infection

We next asked whether TLR priming enables the NAIP/NLRC4 inflammasome to respond to bacteria expressing full-length flagellin containing terminal glutamine-glycine residues in the context of the endogenous flagellar apparatus and T3SS during infection. We generated a *S.* Tm strain in which the terminal arginine at the endogenous FliC locus was replaced with glutamine-glycine. To ensure that NAIP/NLRC4 inflammasome responses to this mutant flagellin were not confounded by FljB, the *Salmonella* alternate phase flagellin, we performed these studies in a *fljB* mutant background. As expected, given normal motility of EPEC as well as other bacteria with flagellins that contain terminal glutamine-glycine rather than arginine residues, *S.* Tm expressing FliC^R475QG^ exhibited wild-type levels of motility (**Fig. S5**). Critically, *S.* Tm carrying a mutation of the FliC C-terminal arginine to glutamine-glycine led to significantly reduced cytotoxicity during infection of WT murine BMDMs, and cytotoxicity was restored by Pam3 priming (**Fig. 5*A***). Furthermore, consistent with our findings that TLR-primed human macrophages are deficient in their ability to respond to immunoevasive flagellins, Pam-primed THP-1 cells had significantly blunted levels of IL-1β release in response to *S.* Tm expressing FliC^R475QG^ (**Fig. 5*B***).

**Figure 5.**
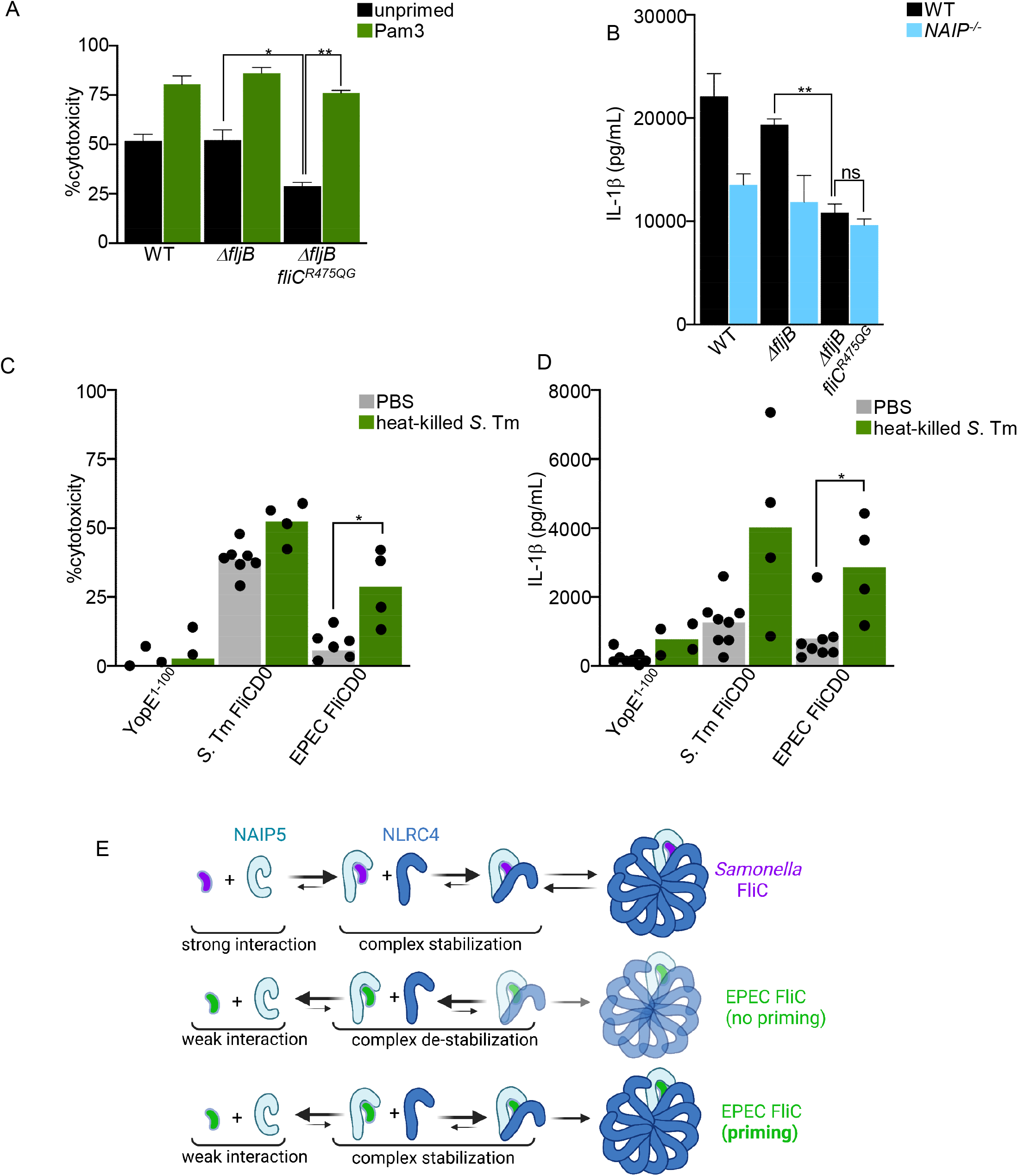
Evasive flagellin differentially impacts murine and human NAIP/NLRC4 inflammasome activation during physiological bacterial infection. (A) WT murine unprimed BMDMs or primed with 0.5 μg/mL Pam3CSK4 for 16 hours were infected with WT, *ΔfljB,* or *ΔfljB fliC^R475QG^ S.* Tm for one hour at an MOI of 20. % cytotoxicity was measured via LDH release. (B) WT or *Naip^-/-^* THP-1 human macrophages were primed with 0.1 μg/mL Pam3CSK4 for 16 hours and infected with WT, *ΔfljB,* or *ΔfljB fliC^R475QG^ S.* Tm for six hours at an MOI of 60. IL-1β release (pg/mL) was measured by ELISA. (C, D) Total peritoneal exudate cells were harvested from WT C57Bl/6J mice intraperitoneally injected with heat-killed *S.* Tm (green bars) or PBS (gray bars) and then infected with the indicated *Yp* strains for 2 hours at an MOI of 20. (C) % cytotoxicity was measured via LDH release. (D) IL-1β release (pg/mL) was measured by ELISA. Each dot represents the mean of one mouse derived from triplicate wells. n=7 for PBS-treated mice and n=4 for heat-killed *S.* Tm-treated mice. Bars represent the means of all mice and statistical analyses were performed using a Mann-Whitney test. (E) Proposed model of how priming enables the NAIP-NLRC4 inflammasome to recognize evasive ligands. (A) Data shown are means ± SEM pooled from three independent experiments and a paired *t* test was performed to assess statistical significance. (B) Data shown are representative of one out of at least three experiments, means ± SEM are shown (n=3) and statistical significance was measured by performing an unpaired *t* test. ns=not significant; *, *P*<0.05, **, *P*<0.01.

During an *in vivo* bacterial infection, the host is naturally exposed to TLR2/4-stimulating ligands. Given that TLR2/4 signaling enables the murine NAIP/NLRC4 inflammasome to sense evasive ligands *in vitro*, we considered that a similar priming of the NAIP/NLRC4 inflammasome would also be expected to occur during *in vivo* exposure to bacteria. To test this model, we primed mice by intraperitoneal injection of heat killed *S.* Tm or PBS control, and isolated total peritoneal cells, which contains macrophages. Intriguingly, peritoneal cells isolated from heat killed *S.* Tm-treated mice but not PBS control mice had increased levels of cytotoxicity and IL-1β release in response to EPEC FliCD0 (**Fig. 5*C-D***). Altogether, these data suggest that priming of the murine NAIP/NLRC4 inflammasome *in vivo* can facilitate cytosolic detection of immunoevasive flagellin.

## Discussion

We report here a T3SS-based system for delivery of T3SS and flagellin-derived NAIP ligands into the host cell cytosol, allowing us to study host responses to these ligands following delivery through their natural route. Our studies reveal that TLR- and p38-mediated upregulation of murine NLRC4 expression enables the NAIP/NLRC4 inflammasome to respond to ligands are currently thought to evade NAIP detection. Interestingly, human macrophages were both basally insensitive to immunoevasive flagellin, and were insensitive to TLR-induced priming, which correlated with their inability to respond to immunoevasive flagellin delivered through the T3SS. Notably, ectopic expression of murine or human NLRC4 in both murine and human macrophages, however, conferred enhanced inflammasome responses against immunoevasive flagellin. Our findings reveal a mechanism for tuning the sensitivity of the NAIP/NLRC4 inflammasome to suboptimal ligands by controlling expression of its adaptor, NLRC4.

In the absence of TLR priming, in addition to the conserved C-terminal leucine residues, which are required for NAIP5-dependent sensing of flagellin (35, 36), the presence of a C-terminal arginine is also required for NAIP5/NLRC4-dependent inflammasome activation via T3SS-delivered FliC (37, 38). Intriguingly, a large number of human-specific bacterial pathogens, including EPEC, *Shigella*, *Helicobacter*, *C. difficile*, among others, produce flagellins that contain C-terminal glutamine, glutamine-glycine, or other non-arginine residues, that empirically do not activate or would be predicted to evade the NAIP5/NLRC4 inflammasome. A number of these flagellins have been reported to not be bound by NAIP5, leading to a model where NAIP5 binding is the determining feature of NAIP5/NLRC4 inflammasome activation (7, 37, 38).

Given that bacterial flagellins and T3SS components that are poor stimulators of the NAIP/NLRC4 inflammasome have been viewed as not binding to their cognate NAIPs, whether a mechanism might exist to enable the host to overcome this type of evasion strategy by flagellins, or other immunoevasive NAIP ligands has not been investigated. Interestingly, while initial reports indicated that *Shigella* flagellin, which contains a terminal -QG, does not bind NAIP5 in yeast two-hybrid based studies (7), it nevertheless leads to NLRC4-dependent lethality when transduced into murine macrophages (36). These data suggest that there are conditions under which the innate immune system can indeed detect evasive flagellins and other NAIP ligands. Our findings demonstrate that TLR-dependent upregulation of NLRC4 expression enables activation of the murine NAIP5/NLRC4 inflammasome in response to evasive flagellins. We find that the main determinant of NAIP5 evasion by flagellin is the presence of a C-terminal glutamine, which can act in a dominant manner in-*cis* to suppress flagellin detection by NAIP5, and that this evasion is specifically overcome by TLR priming. TLR priming of the murine NAIP/NLRC4 inflammasome thus represents a potential adaptation that allows the host to overcome microbial immune evasion in the ongoing evolutionary arms race between pathogens and their hosts.

While the NLRP3 and AIM2 inflammasomes are transcriptionally upregulated in response to inflammatory signals, other than NAIP1, the NAIP/NLRC4 inflammasome is generally not considered to be subject to transcriptional priming (11, 40, 41). NAIP5/NLRC4 inflammasome assembly is initiated by a single NAIP5 subunit that, upon binding flagellin, undergoes a conformational change that recruits and induces a conformational change in the first NLRC4 subunit that in turn uncovers a binding surface to recruit and induce polymerization of subsequent NLRC4 subunits (19, 21, 35, 39, 59). NLRC4 is therefore not thought to be limiting for inflammasome activation. Indeed, our data indicate that TLR priming or ectopic expression of NLRC4 has a limited impact on inflammasome responses to activating flagellin from S. *Tm*, in contrast to evasive flagellin from EPEC. Conversely, blockade of p38 MAPK signaling, which significantly blunted both the Pam3-induced upregulation of *Nlrc4* and the NAIP/NLRC4 response to immunoevasive flagellin, minimally impacted the response to *S*. *Tm* FliCD0. Collectively, these findings indicate that increased expression levels of NLRC4 tune the ability of murine macrophages to detect sub-optimal or immunoevasive flagellins that are poor ligands for NAIP5.

Because EPEC flagellin and other non-activating/evasive ligands were not thought to be bound by their cognate NAIP, they were not thought to be able to nucleate NLRC4 polymerization. We propose that the low affinity between NAIP and evasive ligands results in an unstable complex with a short half-life that does not provide enough time for nucleation and polymerization of NLRC4 (**Fig. 5*E***). However, our findings suggest that cells with elevated NLRC4 expression will exhibit NAIP/NLRC4 avidity, such that a low-affinity ligand-NAIP complex has an increased likelihood to be stabilized by recruitment of the initial NLRC4 subunit, thereby enabling subsequent NLRC4 polymerization and inflammasome activation to take place (**Fig. 5*E***). Our findings imply that in addition to NAIP binding its ligand binding being a key rate-limiting step in NAIP/NLRC4 inflammasome activation, NAIP/NLRC4 avidity may help to set the threshold for inflammasome activation, and suggests that this threshold is tuned by TLR-induced regulation of NLRC4.

Humans express a single NAIP that broadly detects flagellin, T3SS needle, and rod proteins (14, 15, 17). While this promiscuity allows for broader sensing of multiple ligands, human NAIP also does not respond to evasive flagellin or T3SS inner rod proteins. Surprisingly, TLR priming did not induce a response to immunoevasive flagellins in human macrophages, suggesting either that the NAIP/NLRC4 system in humans cannot be primed to sense immunoevasive ligands, or that it may be primed by distinct stimuli. Notably, TLR priming did not actually reduced *NLRC4* expression in human macrophages, in contrast to our observations in mouse macrophages. These findings indicate that the NAIP/NLRC4 inflammasome is subject to distinct regulatory controls in humans and mice. Given the presence of non-activating terminal amino acids in a large number of human specific pathogens (7, 37, 38), including *Shigella*, *Helicobacter*, *Campylobacter*, and the recent findings that NLRC4 serves as an important barrier in mice to colonization by *Shigella* (32, 33), it is tempting to speculate that species-specific transcriptional regulation of the NAIP/NLRC4 inflammasome contribute to the shaping of the repertoire of commensal and pathogenic microbes that colonize specific host niches.

Altogether, our findings uncover a previously unappreciated role for TLR stimulation in transcriptionally regulating the NAIP/NLRC4 inflammasome and licensing the detection of immunoevasive NAIP ligands in murine cells. These findings shed new light on species-specific control of NAIP/NLRC4 inflammasome responses and have implications for the differential ability of pathogens to evade the NAIP/NLRC4 inflammasome in mice and humans.

### Ethics statement

All animal studies were performed in compliance with the federal regulations set forth in the Animal Welfare Act (AWA), the recommendations in the Guide for the Care and Use of Laboratory Animals of the National Institutes of Health, and the guidelines of the University of Pennsylvania Institutional Animal Use and Care Committee. All protocols used in this study were approved by the Institutional Animal Care and Use Committee at the University of Pennsylvania, (Animal Welfare Assurance Number D16-00045/A3079-01, Protocol #804523). All experiments on primary human monocyte-derived macrophages (hMDMs) were performed in compliance with the requirements of the US Department of Health and Human Services and the principles expressed in the Declaration of Helsinki. hMDMs were derived from samples obtained from the University of Pennsylvania Human Immunology Core, and they are considered to be a secondary use of deidentified human specimens and are exempt via Title 55 Part 46, Subpart A of 46.101 (b) of the Code of Federal Regulations.

## Material and Methods

### Bacterial Strains and Growth Conditions

A complete list of strains is found in Table S1. All *Yp* strains were grown in 2x YT broth at 28°C in a shaking incubator at 250 rpm. All *S.* Tm, EPEC and *E. coli* strains were grown in Luria-Bertani Broth (LB) Miller’s formulation at 37°C in a shaking incubator at 250 rpm. When solid medium was required, LB plates containing 1.5% agar supplemented with the appropriate antibiotics were used to incubate either *Yp* for 48hrs or *S.* Tm strains overnight (O/N) at 28°C or 37°C, respectively.

### *Yersinia*-mediated delivery of NAIP ligands

Primers for cloning of NAIP ligands for *Yersinia-*mediated delivery are described in Supplemental Table S2. Briefly, the coding region for the terminal 35 amino acids in the D0 domain of *S.* Tm or EPEC *fliC* genes, *S.* Tm *ssaG*, *ssaI*, and EPEC *escF* was inserted downstream of the *yopE* promoter sequence and the coding sequence of the first 100 amino acids of *yopE* from *Yp* 32777. Constructs were ligated into expression vector pACYC184 using BamHI and SalI restriction sites. Point mutations of the pACYC184 YopE-*S*. Tm or EPEC FliCD0 constructs were generated with the Q5 site-directed mutagenesis kit (New England Biolabs) following manufacturer recommendations. All constructs were validated by Sanger sequencing.

### Isolation and growth of BMDMs

Bone marrow-derived macrophages were isolated and differentiated as previously described (49). Briefly, bone marrow cells isolated from 6-10 week old mice were grown at 37°C, 5% CO_2_ in 30% macrophage media (30% L929 fibroblast supernatant, complete DMEM). BMDMs were harvested in cold PBS on day 7 and replated in 10% macrophage media onto tissue culture (TC)-treated plates prior to subsequent infection. BMDMs were primed with 0.5 μg/mL Pam3CSK4 (Invivogen), 0.05 μg/mL Pam2CSK4 (Invivogen), 100 ng/mL LPS (Sigma), and 100 ng IFN-γ (BioLegend) as indicated in the figure legends for 16-20 hours prior to bacterial infection or harvesting for immunoblot or RNA isolation.

### THP-1 and Human Monocyte-Derived Macrophage growth conditions

THP-1 cells (TIB-202; American Type Culture Collection) were maintained in THP1 growth medium (RPM1 with 10% (vol/vol) heat inactivated FBS + 0.05 nM β-mercaptoethanol + 100 IU/mL penicillin + 100 μg/mL streptomycin) and incubated at 37°C, 5% CO_2_. Human monocytes purified from de-identified healthy human donors were obtained from the University of Pennsylvania Human Immunology Core. To derive the human monocytes into hMDMs, the cells were cultured in RPMI supplemented with 10% (vol/vol) heat inactivated FBS + 2 mM L-glutamine + 100 IU/mL penicillin + 100 μg/mL streptomycin + 50 ng/mL recombinant human M-CSF (Gemini Bio-Products) for 6 days at 37°C, 5% CO_2_. *Nlrc4^-/-^* and *Naip^-/-^* THP-1 cells were previously described (16).

### Infection of macrophages

*Yp* strains were induced to upregulate expression of the T3SS and associated effectors by taking 100 μL of stationary phase overnight cultures and adding to 3 mL 2x YT media containing 20mM sodium oxalate (Sigma) and 20mM MgCl (Sigma), followed by shaking at 28°C for 1 hour and shifting to a 37°C shaking incubator for 2 additional hours. To induce SPI-1 expression in *S.* Tm, 100 μL of overnight cultures were added to 3 mL LB containing 300 mM NaCl and grown at 37°C standing for 3 hours. After induction, the optical density was assessed and used to calculate a multiplicity of infection of 20. Induced cultures were then washed at least three times with pre-warmed BMDM experimental medium (DMEM (high glucose) with L-glutamine + 1 mM Sodium Pyruvate + 1mM HEPES buffer + 10% (vol/vol) heat inactivated FBS + 10% L929 fibroblast cell supernatant). Bacteria were then added to the plated BMDMs and plates were spun down at 1000 rpm for 5 minutes, followed by incubation at 37°C, 5% CO_2_ for one hour, which was then followed by addition of gentamicin (Gold Biotechnology) to a final concentration of 100 μg/mL.

For infection of THP-1-derived human macrophages, two days prior to infection, THP-1 cells were replated in THP-1 growth medium without antibiotics in 48-well plates at a density of 2 x 10^5^ cells per well with 200 nM of phorbol 12-myristate 13-acetate (PMA) for 24 hours to allow differentiation into macrophages. To prime the THP-1 derived macrophages, media was removed 16 hours prior to infection and replaced with THP-1 growth medium containing 100 ng/mL Pam3CSK4. THP-1-derived macrophages were infected at an MOI of 60. For infection of hMDMs, adherent hMDMs were replated in media with 25 ng/mL human M-CSF lacking antibiotics at 1.0 × 10^5^ cells/well in a 48-well plate one day prior to infection and primed with 0.1 μg/mL Pam3CSK4 for 16 hours prior to infection. hMDMs were infected at MOI of 60 as outlined above for the THP-1-derived human macrophages.

### LDH cytotoxicity assays

After infection, cells were spun at 250*g* for 10 minutes to pellet cellular debris. Supernatants were removed and used to assess cytotoxicity via lactate dehydrogenase (LDH) activity. LDH release was quantified using an LDH Cytotoxicity Detection Kit (Roche). Samples were incubated for 25 minutes at room temperature and absorbance at 490 nm was assessed using a spectrophotometer. Percent cytotoxicity was calculated after normalizing to uninfected controls and 100% cell death, which is based on 1% triton X-100-treated cells.

### ELISA

After infection, cells were spun at 250*g* for 10 minutes to pellet cellular debris. Supernatants were removed and used to assess IL-1β levels. For murine IL-1β, an Immulon 2 flat-bottom high binding 96-well plate (BioExpress) was coated with IL-1β capture antibody (eBioscience) at 1 μg/mL in a carbonate-bicarbonate buffer and incubated overnight at 4°C. Next, after washing the plate 5 times with 1x PBS + 0.1 % Tween-20, the plate was blocked in 1% bovine serum albumin (BSA) in 1x PBS for four hours at room temperature. Harvested supernatants from infected cells were diluted 1:4 in 1x PBS and then added to the plate and incubated overnight at 4°C. The plate was then washed six times with 1x PBS + 0.1 % Tween-20 and incubated with biotinylated anti-IL-1β antibody (eBioscience) at 2 μg/mL for two hours at room temperature. After washing six times with 1xPBS + 0.1% Tween-20, the plate was then incubated with streptavidin-HRP (BD Biosciences) for 1 hour at room temperature. The plate was then washed six times with 1xPBS + 0.1% Tween-20, developed with o-phenylenediamine dihydrochloride in citric acid buffer, and signal development stopped with 3M sulfuric acid. Plates were read on a spectrophotometer at 490 nm. IL-1β was quantified based on the standard curve generated using recombinant murine IL-1β (R&D Biosystems). For measuring human IL-1β, an ELISA kit from BD Biosciences was used.

### Immunoblot analysis

To prepare cellular lysate samples for immunoblotting, cells were lysed in 100 μL of lysis buffer (20mM HEPES + 150mM NaCl + 10% (vol/vol) glycerol + 1% (vol/vol) triton X-100 + 1mM EDTA, pH 7.5) with 1x complete protease inhibitor (Roche). NuPAGE LDS sample buffer (4x) (Invitrogen) spiked with 50mM dithiothreitol was added to the lysates, and the samples were boiled at 100°C for 10 minutes. To prepare supernatant samples for immunoblot, supernatants were precipitated with trichloroacetic acid (TCA) overnight. The pelleted TCA samples were washed with acetone, resuspended in TCA-SDS sample buffer (2X SDS buffer + β-mercaptoethanol diluted 1:1 with 0.5M Tris-HCl buffer (pH 8.0) + 4% SDS), and boiled at 100°C for 10 minutes. Lysate and supernatant samples were resolved on NuPAGE 4 to 12% Bis-Tris pre-cast gels (Invitrogen) followed by transfer to 0.2 µm PVDF membrane (EMD Millipore) at 30V for 1 hour using an XCell SureLock Mini-Cell electrophoresis system (Invitrogen). The following primary antibodies were used for immunoblot analysis: 1:5000 anti-β-Actin (Sigma clone AC-74), 1:500 anti-Caspase-1 (Genentech), 1:1000 anti-GSDMD (Abcam [EPR19828]), 1:1000 anti-mouse NLRC4 (Abcam [EPR19733]), 1:1000 anti-human NLRC4 (Cell Signaling Technology, clone D5Y8E). After primary antibody incubation and wash steps, membranes were incubated with species-specific HRP-conjugated secondary antibodies and developed using Pierce ECL Plus (Thermo) or SuperSignal West Femto Maximum Sensitivity Substrate (Thermo) according to the manufacturer’s instructions and visualized on a Bio-rad Gel-Doc XR+ system. Immunoblots were processed and quantified using Image Lab software (Bio-rad).

### Quantitative RT-PCR

RNA was isolated using TRIzol reagent (ThermoFisher) from either 5 x 10^6^ BMDMs or THP-1s following the manufacturer’s protocol. cDNA was prepared from the RNA samples using the high-capacity cDNA reverse transcription kit (Applied Biosystems) per manufacturer’s protocol. Quantitative PCR was conducted with the QuantBio Studio 6 Flex Real-Time PCR system using the PerfeCTa SYBR Green SuperMix (QuantaBio). For analysis, mRNA levels of siRNA-treated cells were normalized to housekeeping gene *GAPDH* (murine) or *HPRT* (human) and fold induction was determined using the 2^−ΔΔCT^ (cycle threshold) method (60). All primers used are listed in Table S3.

### Overexpression of NLRC4 in immortalized macrophages

WT and *Nlrc4^-/-^* murine myeloid progenitors were immortalized using the ER-HoxB8 system (61). To overexpress murine and human NLRC4 in immortalized *Nlrc4^-/-^* murine myeloid progenitors, mscv2.2-IRESGFP retroviral vectors containing murine *Nlrc4* (Addgene plasmid #60199) or human *NLRC4* were used. HEK293T cells were transfected with mscv2.2 vectors containing human or murine *NLRC4* were using lipofectamine 2000 (Invivogen). Briefly, a lipofectamine mixture containing 6 μg of pVSV-G + 12 μg of pCL-ECO + 20 μg of mscv2.2 vector was transfected per each 10-cm plate of HEK293T cells. Retroviral supernatants were filtered and concentrated with Lenti-X concentrator (Takara Bio) according to manufacturer’s instructions. *Nlrc4^-/-^* immortalized myeloid progenitors were spinfected on fibronectin-coated (10 μg/mL) 12-well plates with the concentrated retrovirus in the presence of 25 μg/mL polybrene at 1750 rpm for 90 minutes. After spinfection, fresh R10 media containing 0.5 μM estrogen and 10 ng/mL GM-CSF was added to the cells. After 48 hours, the same cells were spinfected again with more retroviral supernatant. The transduced progenitor cells were allowed to recover and expand over the course of three weeks. Successfully transduced cells were sorted by flow cytometry assisted cell sorting based on expression of GFP. Expression of murine or human NLRC4 in the immortalized myeloid progenitors was verified via immunoblot prior to use. THP-1 human monocyte-derived macrophages transduced with the MigR1 retroviral vector alone or overexpressing *hNLRC4* were previously described (62).

### Generation of S. Tm *Δfljb::kan fliC^R475QG^*

To mutate SL1344 to express flagellin terminating in a glutamine-glycine in place of arginine, allelic exchange using the suicide vector pCVD442 was performed. Briefly, mutagenic primers were designed to span 800 base pairs upstream and downstream of the C-terminal portion *flic* gene harboring the arginine to glutamine-glycine mutation. This construct was then ligated into pCVD442 using SacI and SalI restriction sites and transformed into DH5α λpir and resulting transformants were sequence verified to carry mutation of interest. Next, a triparental mating using the donor pCVD442 *fliC^R475QG^* DH5α λpir strain, a helper *E. coli* strain, and recipient SL1344 *Δfljb::kan* was performed. These strains were grown O/N and supplemented with antibiotics when appropriate. In a microcentrifuge tube, 400 μL of the donor, 400 μL of the helper and 600 μL of the recipient were mixed and washed in LB, and then spot plated onto LB agar. After 6hrs of incubation at room temperature (RT), this conjugation mixture was harvested, and serially diluted in sterile PBS and plated onto LB agar plates containing Strep and Amp. After incubating O/N, single colonies from the conjugation plate were inoculated into LB and grown for 3 hours until an OD between 0.8-0.9 was reached. These cultures were serially diluted and then plated on LB agar plates containing 20% sucrose for sucrose counterselection and allelic exchange. The resulting mutant strain *S.* Tm Δ*fljb::kan fliC^R475QG^* was sequence-verified to verify that the desired mutation was present.

### Ex vivo infections

Two days after intraperitoneal injection of 10- to 12-week-old C57BL/6J male and/or female mice with 2 x10^7^ CFU of heat killed *S.* Tm or PBS, peritoneal cells were isolated by peritoneal lavage with 7 mL of sterile PBS. Samples were spun for 5 minutes at 1500 rpm at 4°C. Cells were then resuspended in DMEM with 10 % (vol/vol) FBS and washed a total of three times. 1 x 10^5^ total peritoneal cells per well were plated in 80 μL of DMEM with 10 % (vol/vol) FBS and incubated for 3-5 hours to let the cells adhere to the plate. Prior to infecting with bacteria, the peritoneal cells were washed one time with DMEM containing 10 % (vol/vol) FBS.

### Statistical analysis

Graphing and statistical analyses of data were performed using Prism 9 software (GraphPad). Statistical significance was determined using the statistical tests indicated in each figure legend. Differences were considered statistically significant if the *P* value was less than or equal to 0.05.

## Acknowledgments

We would like to thank Dr. Thirumala-Devi Kanneganti (St. Jude Children’s Research Hospital) for *Naip5^-/-^* bone marrow, Dr. Dieter Schifferli (University of Pennsylvania) for the EPEC strain, Dr. Annelise Snyder for initial flagellin constructs, members of the Shin and Brodsky labs for scientific discussion, Dr. Alex Hoffmann (UCLA) for scientific discussion, and Dr. Patrick Mitchell (University of Washington Seattle) for scientific discussion and critical reading. These studies were supported by National Institutes of Health grants AI128520, AI139102, AI163596 (IEB), AI123243, AI118861, AI151476 (SS), F32AI164655 (JPG), and R01HD098428 (SC), National Science Foundation Graduate Research Fellowship DGE-1321851 (MSE), the Mark Foundation Center for Radiobiology and Immunology-Project 4 (IEB), and Investigators in the Pathogenesis of Infectious Disease Awards from the Burroughs Wellcome Fund (SS and IEB).

