## Supplemental Information for "TLR priming licenses NAIP inflammasome activation by immunoevasive ligands"

### Supporting Information for
***TLR priming licenses NAIP inflammasome activation by immunoevasive ligands***

James P. Grayczyk[1,†], Marisa S. Egan[2], Luying Liu[1], Emily Aunins[1], Meghan A. Wynosky-Dolfi[3,††], Scott Canna[4], Andy J. Minn[5, 6, 7, 8], Sunny Shin[2], and Igor E. Brodsky[1]

[1]Department of Pathobiology, University of Pennsylvania School of Veterinary Medicine, Philadelphia, PA, USA
[2]Department of Microbiology, University of Pennsylvania Perelman School of Medicine, Philadelphia, PA, USA
[3]Department of Pathobiology, University of Pennsylvania School of Veterinary Medicine, Philadelphia, PA, USA
[4]Department of Pediatrics, Division of Rheumatology, Children's Hospital of Philadelphia, Philadelphia, PA
[5]Department of Radiation Oncology, [6]Abramson Family Cancer Research Institute, [7]Parker Institute for Cancer Immunotherapy, [8]Mark Foundation Center for Immunotherapy, Immune Signaling, and Radiation, University of Pennsylvania Perelman School of Medicine, Philadelphia, PA, USA

[†]Current address: Oncology Discovery, AbbVie Inc., 1 North Waukegan Rd., North Chicago, IL 60064
[††]Current address: Immunology Research Unit, GlaxoSmithKline, Collegeville PA

**This PDF file includes:**

Figures S1 to S5
Tables S1 to S3
SI References

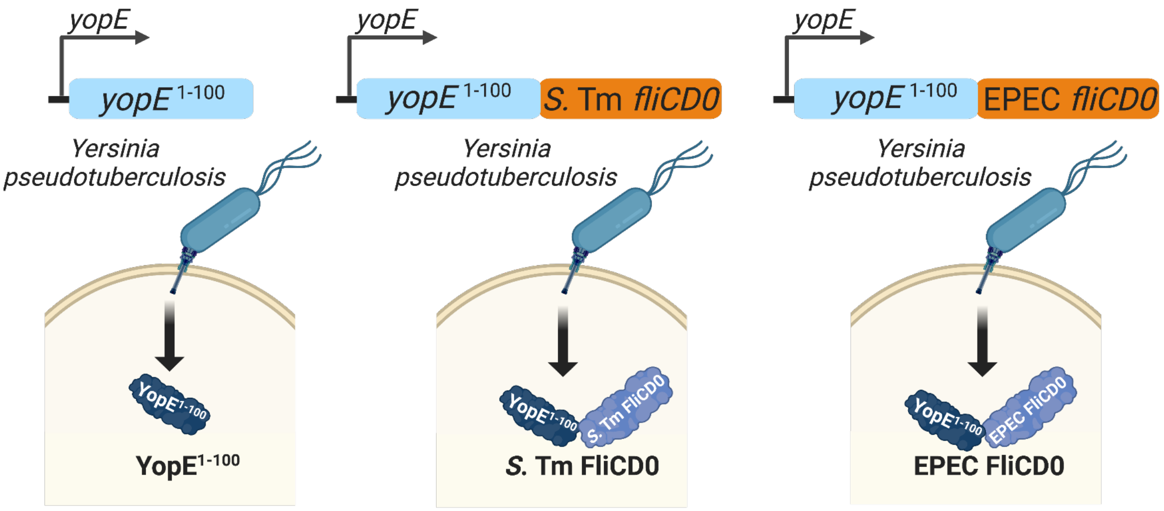


**Fig. S1**. Design of *Yersinia pseudotuberculosis* T3SS ligand delivery system. A heterologous delivery system using the T3SS of the enteric gram-negative pathogen *Yersinia pseudotuberculosis* (*Yp*) to deliver NAIP ligands was made by fusing the C-terminal FliCD0 region to the N-terminal 100 amino acids of the T3SS-secreted *Yersinia* outer protein E ($YopE^{1\text{-}100}$). This fusion protein of $YopE^{1\text{-}100}$ with *S.* Tm or EPEC FliCD0 is under the control of the native *yopE* promoter.

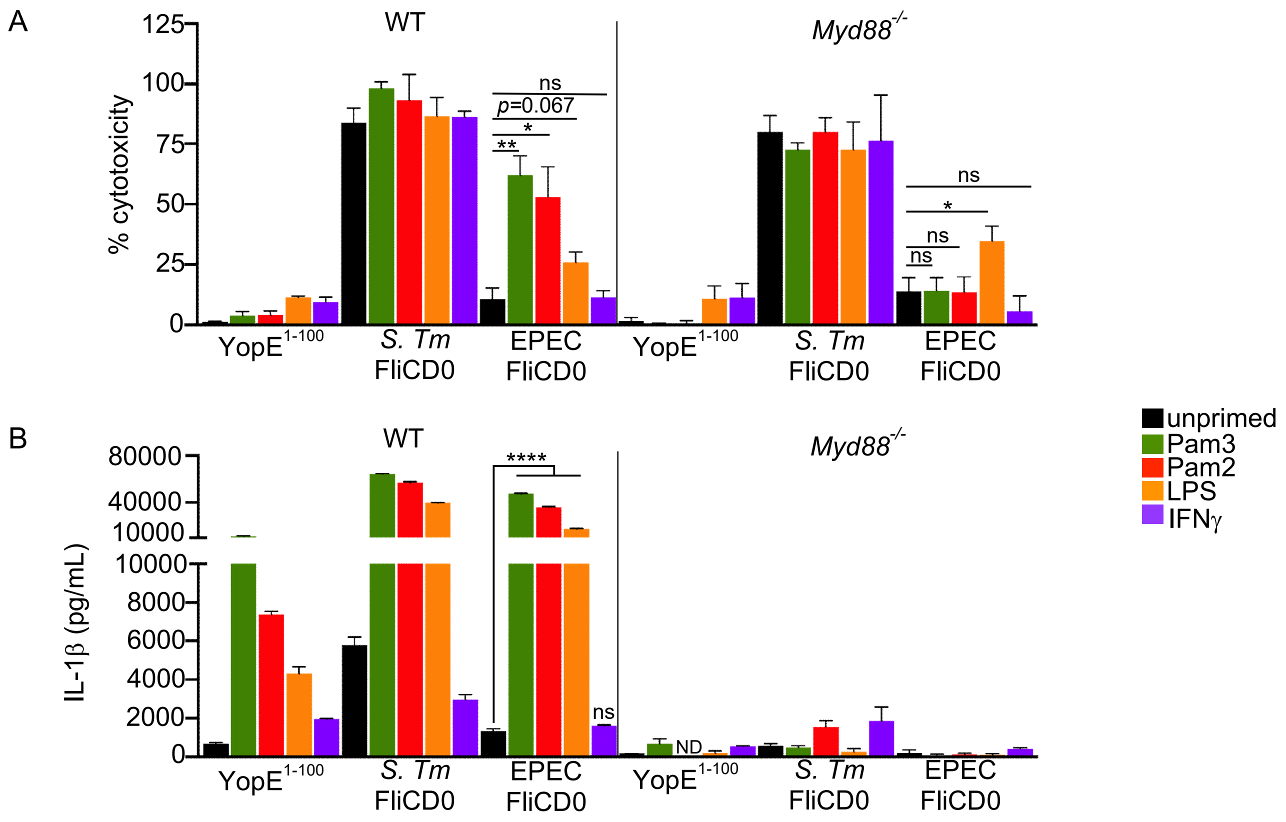


**Fig. S2.** TLR2 and TLR4 priming enhance the inflammasome response to EPEC FliCD0 in mouse macrophages. (A,B) WT murine unprimed BMDMs or primed with 0.5 μg/mL Pam3CSK4, 0.05 μg/mL Pam2CSK4, 100ng/mL LPS or 100ng IFN-γ for 16 hours were infected with the indicated strains for four hours at an MOI of 20. (A) % cytotoxicity was measured via lactase dehydrogenase (LDH) release. (B) IL-1β release (pg/mL) was measured by ELISA. (A) Data shown are means ± SEM pooled from three independent experiments for unprimed, Pam3CSK4-primed, Pam2CSK4-primed, or LPS-primed conditions or two independent experiments for IFN-γ-primed condition. A paired *t* test was performed to assess statistical significance. (B) Data shown are representative of three independent experiments. Combined means ± SEM are shown (n=3). Statistical significance was measured by performing an unpaired *t* test. ND=not detected; ns=not significant; *, $P<0.05$, **, $P<0.01$,***, $P<0.001$,****, $P<0.0001$.

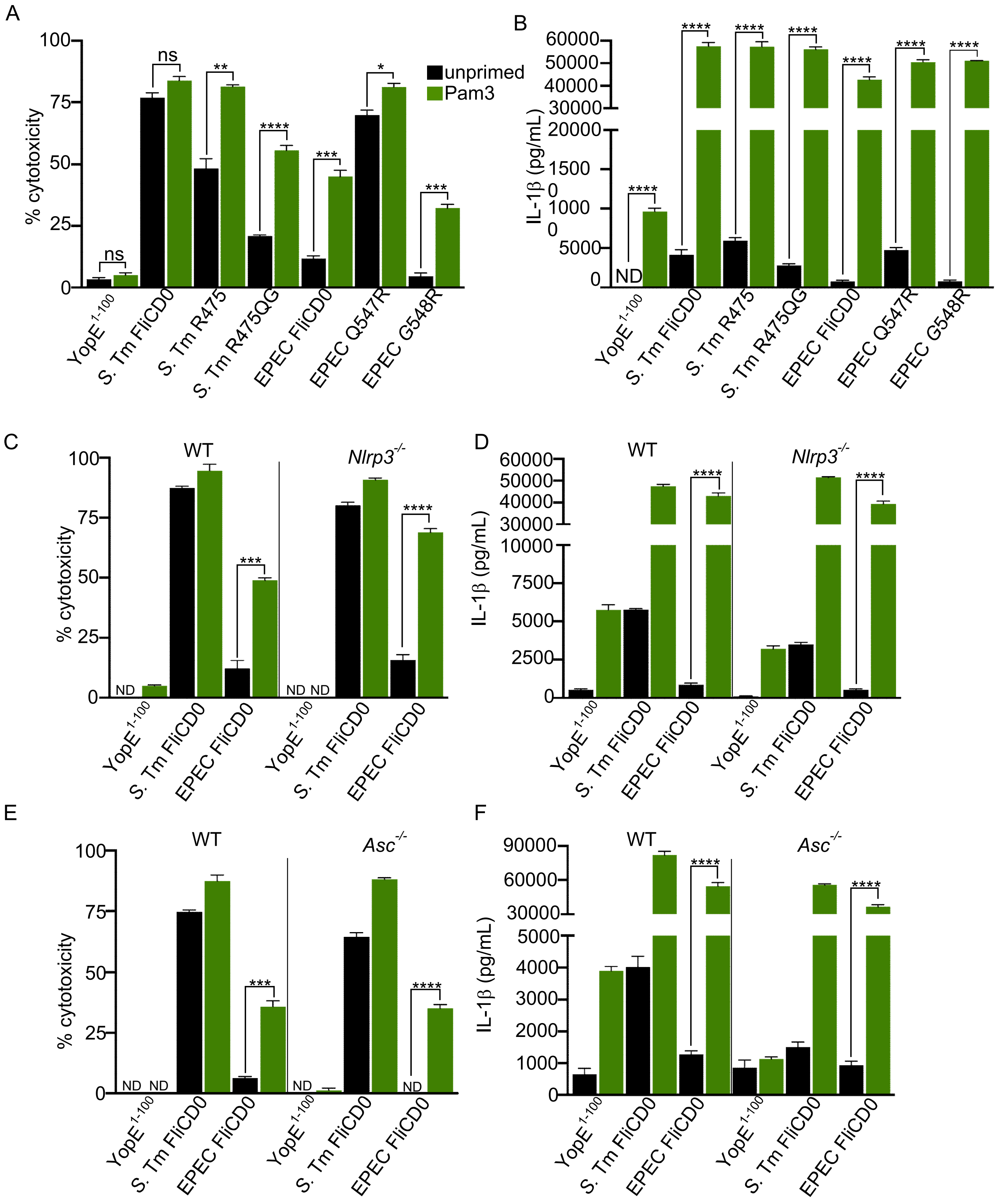
A
B
C
D
E
F
% cytotoxicity
IL-1β (pg/mL)
unprimed
Pam3
ns
ND
WT
Nlrp3-/-
Asc-/-
YopE1-100
S. Tm FliCD0
S. Tm R475
S. Tm R475QG
EPEC FliCD0
EPEC Q547R
EPEC G548R

**Fig. S3.** TLR-mediated detection of EPEC FliCD0 by the NAIP/NLRC4 inflammasome is independent of NLRP3 and ASC. (A-F) WT, *Myd88*$^{-/-}$, *Nlrp3*$^{-/-}$, or *Asc*$^{-/-}$ murine unprimed BMDMs or primed with 0.5 μg/mL Pam3CSK4 for 16 hours were infected with the indicated strains for four hours at an MOI of 20. (A, C, E) % cytotoxicity was measured via LDH release. (B, D, F) IL-1β release (pg/mL) was measured by ELISA. (A, C, E) Data shown are means ± SEM pooled from three independent experiments for unprimed, Pam3CSK4-primed, Pam2CSK4-primed, or LPS-primed conditions or two independent experiments for IFN-γ-primed conditions. A paired *t* test was performed to assess statistical significance. (B, D, F) Data shown are representative of three independent experiments, means ± SEM are shown (n=3) and statistical significance was measured by performing an unpaired *t* test. ND= not detected; ns=not significant; *, $P<0.05$, **, $P<0.01$,***, $P<0.001$,****, $P<0.0001$

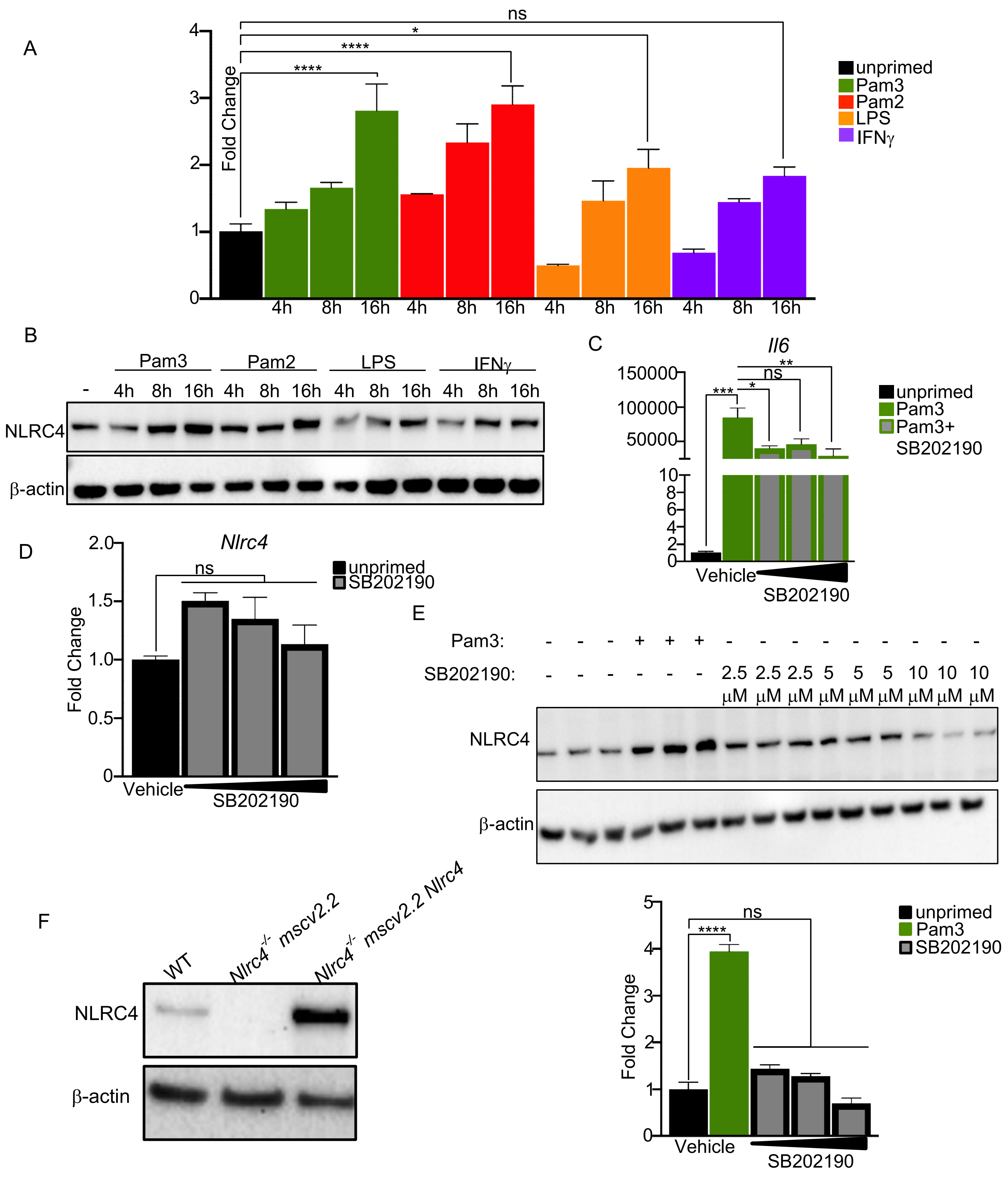
A
ns
*
****
****
Fold Change
0
1
2
3
4
4h 8h 16h 4h 8h 16h 4h 8h 16h 4h 8h 16h
unprimed
Pam3
Pam2
LPS
IFNγ
B
Pam3
Pam2
LPS
IFNγ
- 4h 8h 16h 4h 8h 16h 4h 8h 16h 4h 8h 16h
NLRC4
β-actin
C
Il6
150000
100000
50000
10
8
6
4
2
0
***
*
ns
**
unprimed
Pam3
Pam3+ SB202190
Vehicle
SB202190
D
Nlrc4
2.0
1.5
1.0
0.5
0
Fold Change
ns
unprimed
SB202190
Vehicle
SB202190
E
Pam3: - - - + + + - - - - - - - - -
SB202190: - - - - - - 2.5 µM 2.5 µM 2.5 µM 5 µM 5 µM 5 µM 10 µM 10 µM 10 µM
NLRC4
β-actin
F
WT
Nlrc4-/- mscv2.2
Nlrc4-/- mscv2.2 Nlrc4
NLRC4
β-actin
5
4
3
2
1
0
Fold Change
****
ns
unprimed
Pam3
SB202190
Vehicle
SB202190

**Fig. S4.** TLR priming but not IFN-$\gamma$ increases NLRC4 transcript and protein levels in macrophages. (A, B) WT murine BMDMs were unprimed or primed with 0.5 $\mu$g/mL Pam3CSK4, 0.05 $\mu$g/mL Pam2CSK4, 100ng/mL LPS, or 100ng IFN-$\gamma$ for 4, 8, or 16 hours. (A) *Nlrc4* transcript level relative to *Gapdh* was assessed via qRT-PCR. Fold change in *Nlrc4* transcript levels was analyzed relative to unstimulated control. (B) NLRC4 protein levels were analyzed via immunoblot. Data representative of three independent experiments. (C-E) WT murine BMDMs were unprimed or primed with 0.5 μg/mL Pam3CSK4. One hour prior to priming, BMDMs were treated with 2.5 μM, 5 μM, or 10 μM p38 inhibitor SB202190 or DMSO vehicle control. (C, D) *Il6* or *Nlrc4* transcript levels relative to *Gapdh* was assessed via qRT-PCR. Fold change was analyzed as relative to unstimulated control. (E) NLRC4 protein levels were analyzed via immunoblot. NLRC4 protein intensity was quantified relative to $\beta$–actin loading controls, and fold change was assessed relative to unstimulated DMSO treated cells. Data representative of three independent experiments. (F) NLRC4 protein levels in WT or *Nlrc4*$^{-/-}$ mscv2.2-transduced ER-HoxB8 macrophages were assessed via immunoblot. Data representative of two independent experiments. (A, C-E) Data representative of three independent experiments. Means ± SEM are shown (n=3) and statistical significance was measured by performing a 1-way ANOVA (C-E). ns=not significant; *, *P*<0.05, **, *P*<0.01,***, *P*<0.001,****, *P*<0.0001

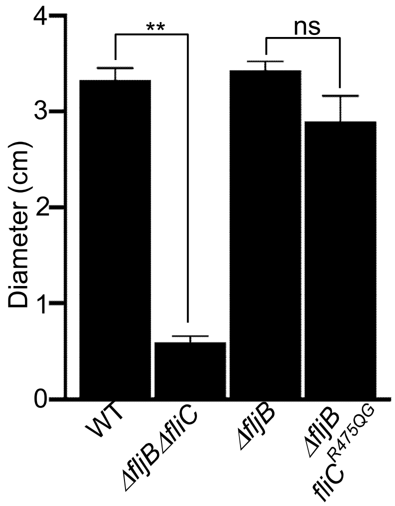


**Fig. S5. *S.* Tm *ΔfljB fliC*$^{R475QG}$ mutant does not have impaired flagellar-based motility.** LB plates containing 0.4% agar were stabbed with optical density-normalized overnight cultures of WT, *ΔfljBfliC, ΔfljB,* or *ΔfljB fliC*$^{R475QG}$ *S.* Tm. After 12-16 hours incubation at 37°C, the diameter of the colony was measured to quantify motility. Data shown are pooled from 3 independent experiments, mean ± SEM are shown, and statistical significance was measured by performing a paired *t* test. ns=not significant; **, *P*<0.01.

**Table S1**. List of strains.

| Strain | Description | Source |
|---|---|---|
| *Yersinia pseudotuberculosis* (*Yp)* | strain 32777, serogroup O1 | (1) |
| *ΔyopJ Yp* | strain 32777 harboring a clean deletion of *yopJ* (or is it the point mutant?) | (2) |
| *Salmonella enterica* serovar Typhimurium | SL1344 | (3) |
| *Escherichia coli* DH5α | Cloning strain | - |
| *Escherichia coli* DH5α λpir | Cloning strain for R6K plasmid replication | - |
| *Escherichia coli* CR19 | Conjugation helper strain | (4) |
| Enteropathogenic *Escherichia coli* | E2348 | Dieter Schifferli |
| *ΔyopJ Yp* pACYC184 yopE alone | strain 32777 with a mutated *yopJ* with the YopE promoter sequence with the first 100bp of yopE expressed in pACYC184 | This study |
| *ΔyopJ Yp* pACYC184 yopE-*S.* Tm FliCD0 | strain 32777 with a mutated *yopJ* with the YopE promoter sequence with the first 100bp of yopE fused to *S.* Tm FliCD0 expressed in pACYC184 | This study |
| *ΔyopJ Yp* pACYC184 yopE-EPEC FliCD0 | strain 32777 with a mutated *yopJ* with the YopE promoter sequence with the first 100bp of yopE fused to EPEC FliCD0 expressed in pACYC184 | This study |
| *Yp* pACYC184 yopE alone | strain 32777 with the YopE promoter sequence with the first 100bp of yopE expressed in pACYC184 | This study |
| *ΔyopJ Yp* pACYC184 yopE-*S.* Tm FliCD0 | strain 32777 with the YopE promoter sequence with the first 100bp of yopE fused to *S.* Tm FliCD0 expressed in pACYC184 | This study |
| *ΔyopJ Yp* pACYC184 yopE-EPEC FliCD0 | strain 32777 with the YopE promoter sequence with the first 100bp of yopE fused to EPEC FliCD0 expressed in pACYC184 | This study |
| *ΔyopJ Yp* pACYC184 yopE-*S.* Tm FliCD0 R475 | strain 32777 with a mutated *yopJ* with the YopE promoter sequence with the first 100bp of yopE fused to *S.* Tm FliCD0 R475 expressed in pACYC184 | This study |
| *ΔyopJ Yp* pACYC184 yopE-*S.* Tm FliCD0 R475QG | strain 32777 with a mutated *yopJ* with the YopE promoter sequence with the first 100bp of yopE fused to *S.* Tm FliCD0 R475QG expressed in pACYC184 | This study |
| *ΔyopJ Yp* pACYC184 yopE-EPEC FliCD0 Q547R | strain 32777 with a mutated *yopJ* with the YopE promoter sequence with the first 100bp of yopE fused to EPEC FliCD0 Q547R expressed in pACYC184 | This study |
| *ΔyopJ Yp* pACYC184 yopE-EPEC FliCD0 G548R | strain 32777 with a mutated *yopJ* with the YopE promoter sequence with the first 100bp of yopE fused to EPEC FliCD0 G548R expressed in pACYC184 | This study |
| *ΔyopJ Yp* pACYC184 yopE-*S.* Tm SsaG | strain 32777 with the YopE promoter sequence with the first 100bp of yopE fused to *S.* Tm *ssaG* expressed in | This study |

| | | |
|---|---|---|
| | pACYC184 | |
| *ΔyopJ Yp* pACYC184 yopE-*S.* Tm SsaI | strain 32777 with the YopE promoter sequence with the first 100bp of yopE fused to *S.* Tm *ssaI* expressed in pACYC184 | This study |
| *ΔyopJ Yp* pACYC184 yopE-EPEC EscI | strain 32777 with the YopE promoter sequence with the first 100bp of yopE fused to EPEC *EscI* expressed in pACYC184 | This study |
| *Salmonella enterica* serovar Typhimurium *Δfljb::kan* | SL1344 with a kanamycin resistance cassette in place of *fljb* | (5) |
| *Salmonella enterica* serovar Typhimurium *Δfljb::kan flic*$^{R475QG}$ | SL1344 with a kanamycin resistance cassette in place of *fljb* and *fliC*$^{R475QG}$ | This study |
| *Salmonella enterica* serovar Typhimurium *ΔfljbΔflic* | SL1344 harboring a clean deletion of *fljb* and *flic* | This study |

**Table S2.** Primer list for bacterial strains.

| Name | Sequence | Purpose |
|---|---|---|
| YopEFliCD0For | CACAAATGCCAAGTCCTACGAGCCGTATCGAAGATTCCGA | YopE *S.* Tm FliCD0 |
| YopEFliCD0Rev | CGTAGGACTTGGCATTTGTG | YopE *S.* Tm FliCD0 |
| YopEFliCepecA | TAAGCGGATCCCCAACTTTGACACCGATA | YopE EPEC FliCD0 and YopE alone |
| YopEFliCepecB | AATACGGGACTGCGCCGTAGGACTTGGCATT | YopE EPEC FliCD0 |
| YopEFliCepecC | AATGCCAAGTCCTACGGCGCAGTCCCGTATT | YopE EPEC FliCD0 |
| YopEFliCepecD | TAAGCGTCGACTTAACCCTGCAGCAGAG | YopE EPEC FliCD0 |
| YopEnegB | TAAGCGTCGACCGTAGGACTTGGCATT | YopE$^{1-100}$ |
| SL1344LL_Q5F | ATAAGTCGACCGATGCCCTTGA | *S.* Tm FliCD0 R475 |
| SL1344LL_Q5R | CACAGTAAAGAGAGGACGTTTTGCG | *S.* Tm FliCD0 R475 |
| SL1344QG_Q5F | GGATAGGACCGATGCCCTTGAGAGC | *S.* Tm FliCD0 R475QG |
| SL1344QG_Q5R | CTGCAGTAAAGAGAGGACGTTTTGCG | *S.* Tm FliCD0 R475QG |
| EPECQ5F | TCTCTGCTGCGGTGATAAGTCGACCG | EPEC FliCD0 Q547R |
| EPECQ5R | CAGAACCTGCTGCGGTAC | EPEC FliCD0 Q547R |
| EPECQR_Q5F | TTAAGTCGACCGATGCCCTTGA | EPEC FliCD0 G548R |
| EPECQR_Q5R | CGCTGCAGCAGAGACAGAACCTGC | EPEC FliCD0 G548R |
| P1OEQG | TAAGCAGAGCTCctggaaaatattacgccaaagttac | *flic::R475QG* |
| P2OEQG | GCAGTAAAGAGCTATCCCTAATCAATCGCC | *flic::R475QG* |
| P3OEQG | CTCTTTACTGCAGGGATAGGGCGATTGATT | *flic::R475QG* |
| P4OEQG | TAAGCAGTCGACAGCGAGGTTTTTACCTTGCTATC | *flic::R475QG* |

**Table S3.** Primer list for qPCR.

| Gene | Sequence | Source |
|---|---|---|
| Mouse *Nlrc4* F | CAGGTGGTCTGATTGACAGC | (6) |
| Mouse *Nlrc4* R | CCCCAATGTCAGACAAATGA | |
| Mouse *Naip1* F | TGCCCAGTATATCCAAGGCTA | |
| Mouse *Naip1* R | AGACGCTGTCGTTGCAGTAAG | |
| Mouse *Naip2* F | TTTTGTGAATCCCTGGGTCA | |
| Mouse *Naip2* R | TGTAGAAAAGGCCTGCTTTGA | |
| Mouse *Naip5* F | AAGGAGATGACCCCTGGAAG | |
| Mouse *Naip5* R | TGACCCAGGACTTCACAAAA | |
| Mouse *Naip6* F | TTTTGTGAAGTCCTGGGTCAG | |
| Mouse *Naip6* R | CAATGTCCTTTTTGCCAGTG | |
| Mouse *Il6* F | TCAATTCCAGAAACCGCTATGAAG | (7) |
| Mouse *Il6* R | CGTTGTTCATACAATCAGAATTGCCA | |
| Mouse *Gapdh* F | CTCCCACTCTTCCACCTTCG | (8) |
| Mouse *Gapdh* R | CCACCACCCTGTTGCTGTAG | |
| Human *NLRC4* F | CATAGTCAAGTCTCTGTCAAGTGAACCCTGT | (9, 10) |
| Human *NLRC4* R | GCTGTTCTAGCACGTTCATCCTGTCG | |
| Human *NAIP* F | AAGCATCCGCCCAGCTCTTGA | |
| Human *NAIP* R | TATTGCCCTCCAGATCCACAGACAGTTC | |
| Human *IL6* F | ACTCACCTCTTCAGAACGAATTG | |
| Human *IL6* R | CCATCTTTGGAAGGTTCAGGTTG | |
| Human *HPRT1* F | CCTGGCGTCGTGATTAGTGAT | (11) |
| Human *HPRT1* R | AGACGTTCAGTCCTGTCCATAA | |
